# Bacterial lipopolysaccharide induces settlement and metamorphosis in a marine larva

**DOI:** 10.1101/851519

**Authors:** Marnie L. Freckelton, Brian T. Nedved, You-Sheng Cai, Shugeng Cao, Helen Turano, Rosanna A. Alegado, Michael G. Hadfield

## Abstract

How larvae of the many phyla of marine invertebrates find places appropriate for settlement, metamorphosis, growth and reproduction is an enduring question in marine science. Biofilm induced metamorphosis has been observed in marine invertebrate larvae from nearly every major marine phylum. Despite the widespread nature of this phenomenon the mechanism of induction remains poorly understood. The serpulid polychaete *Hydroides elegans* is a well-established model for investigating bacteria-induced larval development. A broad range of biofilm bacterial species elicit larval metamorphosis in *H. elegans* via at least two mechanisms, including outer membrane vesicles and phage-tail bacteriocins. We investigated the interaction between larvae of *H. elegans* and the inductive bacterium *Cellulophaga lytica*, which produces an abundance of OMVs but not phage-tail bacteriocins. We asked whether the OMVs of *C. lytica* induce larval settlement due to cell membrane components or through delivery of specific cargo. Employing a biochemical structure-function approach with a strong ecological focus, the cells and outer membrane vesicles produced by *C. lytica* were interrogated to determine the nature of the inductive molecule. Here we report that the cue produced by *C. lytica* that induces larvae of *H. elegans* to metamorphose is lipopolysaccharide (LPS). The widespread prevalence of LPS and its associated taxonomic and structural variability suggest it may be a broadly employed cue for bacterially induced larval settlement of marine invertebrates.

**Significance Statement:** New surfaces in the sea are quickly populated by dense communities of invertebrate animals, whose establishment and maintenance require site-specific settlement of larvae from the plankton. Although it is recognized that larvae selectively settle in sites where they can metamorphose and thrive, and that the biofilm bacteria residing on these surfaces supply inductive cues, the nature of the cues used to identify ‘right places’ has remained enigmatic. In this paper, we reveal that lipopolysaccharide (LPS) from the outer membrane of a marine Gram-negative bacterium cue metamorphosis for a marine worm and discuss the likelihood that LPS provides the variation necessary to explain settlement site selectivity for many of the bottom-living invertebrate animals that metamorphose in response to bacterial biofilms.

## Introduction

More than two-thirds of the earth’s surface is covered by the oceans. The solid surface beneath these salt waters is greater than that of all the jungles, rain forests, boreal forests, prairies, desserts, tundras, and mountains combined. Yet, we know more about the dynamics of the biological world living above the ocean’s shores than we do about the enormously diverse benthic communities that extend from the higher reaches of the intertidal to the greatest depths of the seas. With global climate change causing massive melting of the earth’s glaciers, sea levels are rising, and the amount of land covered by the seas is increasing dramatically. The establishment and maintenance of benthic marine animal communities depends heavily on recruitment of minute larvae from the plankton. Larval recruitment is an essential process not only for all marine benthic communities, but also for creating harvestable populations of mariculture species such as clams, oysters, and shrimp. It is also the initial process in the establishment of costly biofouling communities on the hulls of ships and in the pipes that bring cooling seawater to coastal electrical generators (1). A critical element of this process that remains poorly understood is how larvae select habitats suitable not just for attachment and metamorphosis, but long-term survival, growth and reproduction (2, 3).

In recent decades we have learned that the cues most invertebrate larvae employ to select right places to metamorphose are almost universally biological, the majority originating from microbes residing in surface biofilms (4, 5). Biofilm- or bacterial-induced settlement has been shown for larvae of sponges (6–8), cnidarians (9, 10), bryozoans (11, 12), molluscs (13–16), annelids (17), echinoderms (18, 19), crustaceans (20, 21) and urochordates (22), i.e., all of the major invertebrate phyla. Although the number of larval types recorded to settle and metamorphose in response to bacterial films is now so great as to suggest a nearly universal mechanism for both settlement induction and larval response, we remain largely ignorant about the diversity of bacteria that stimulate larvae to settle, the chemical nature of the bacterial cues, and the mechanisms involved in larval recognition of bacteria.

Larvae of the serpulid polychaete *Hydroides elegans* provide an excellent model for investigating bacteria-stimulated settlement (17, 23). *H. elegans* is a cosmopolitan member of the benthic and biofouling communities in warm-water bays and ports throughout the world (23). As is the case for a vast majority of marine invertebrates, an external stimulant induces an extremely rapid series of morphogenetic events that quickly transform the swimming larva of *H. elegans* into a bottom-living, tube-dwelling juvenile that will grow rapidly into a reproductive adult (23). Larvae of *H. elegans* typically do not settle in the absence of a biofilm, and this response follows direct contact with surface biofilms (17, 23–28). Investigations of the complex biofilm communities that induce settlement and metamorphosis in *H. elegans* and the cues that they produce must take place in the context of this ecology. Inductive cues must be naturally entrained within the biofilm and fast acting, because even in a low turnover environment such as Pearl Harbor, Hawai□i, current speeds and turbulence at the biofilm interface would otherwise rapidly disperse and remove dissolved compounds (29–31).

Evidence has accumulated that it is particular bacterial species residing in the biofilms that induce the metamorphic response in the larvae of *H. elegans* (5, 28, 32). Inductive bacteria from a broad range of phyla, including both Gram-negative and Gram-positive strains, are inductive when cultured in mono-species biofilms (24, 28, 32–35). However, most biofilm-bacterial species do not induce settlement in *H. elegans* (28, 36). To date, we have identified at least two different induction mechanisms from two different Gram-negative bacteria, *Pseudoalteromonas luteoviolacea* and *Cellulophaga lytica.* For *P. luteoviolacea* strain HI1, a highly inductive biofilm species isolated from Pearl Harbor, Hawai□i and genetically characterized in our lab, we identified specific structural elements derived from phage-tail proteins that are in some manner involved in settlement induction (37–39). However, examination of other inductive bacterial species revealed that induction of larval settlement cannot alone be explained by phage-tail bacteriocins. *Cellulophaga lytica,* another biofilm bacterium, induces larvae of *H. elegans* to settle by an entirely different mechanism. Although cell-free preparations from broth cultures of *C. lytica* induce the tubeworm larvae to metamorphose, analysis of the genome of *C. lytica* (40) yielded none of the genes that transcribe the phage-tail elements of bacteriocins (34). Additionally, transmission electron microscopy (TEM) of inductive supernatants from *C. lytica* revealed an abundance of outer membrane vesicles (OMVs) and none of the bacteriocins found in similar preparations from *P. luteoviolacea* (34).

We have previously suggested that such membrane vesicles should be considered for their potential to be a common mechanism of interaction between biofilm bacteria and invertebrate larvae (35). Gram-negative bacteria ubiquitously produce OMVs (41, 42), but their structures and contents are variable per species and strain. Interactions between microbes and eukaryotes are mediated largely by a suite of conserved molecules knowns as Microbe-Associated Molecular Patterns (MAMPs) (43, 44). Two major MAMP classes, lipopolysaccharide and peptidoglycan, comprise important components of the envelope of both whole cells and OMVs. Additionally, OMVs may enclose genetic material, virulence factors, proteins, small bioactive metabolites and lipids (41). Consequently, OMVs provide the mechanism for multiple ecological roles fulfilled by Gram-negative bacteria, including cell-to-cell signaling and transfer of DNA, proteins and small signaling molecules between cells (41, 42, 45–53). The ubiquity of OMVs means that the response to their detection by a eukaryote is entirely context dependent. In the present study, we asked whether the OMVs of *C. lytica* induce larval settlement because they are accessible pieces of bacterial cell membranes or because they deliver a specific cargo.

To gain key insight into the molecular structure of the larval settlement inducer produced by *C. lytica* and to determine if it is a protein, nucleic acid, or lipid, we subjected the OMV fractions to a battery of broad-spectrum enzymatic and chemical treatments and then tested the treated OMVs for loss of larval settlement-inducing activity. Subsequently, because OMVs are composed of pieces of the bacterial outer membrane and because of the need for larger quantities of compounds for treatments and assays, the components of the cell envelope (whole membrane, peptidoglycan and lipopolysaccharide) were assayed directly by extraction from whole cells. These approaches, including *in silico* analyses of the bacterial genome, HPLC fractionation of methanol extracts, and enzymatic destructions of OMVs and isolated cell membrane components, combined with larval settlement assays, allowed us to reject all but bacterial LPS as cues for larval settlement. As the outermost element of Gram-negative bacterial cells, membrane LPS provides the surface-contact stimulus indicated by behavioral studies of larval settlement in *H. elegans* (26). We conclude LPS is a key element in selective larval settlement by *H. elegans*, and we predict this will hold true for many other benthic marine invertebrate animals.

## Results

### OMV production, size, and metamorphic bioactivity as a function of growth phase in *C. lytica*

OMVs isolated from broth cultures of *C. lytica* induced metamorphosis in larvae of *H. elegans* at all growth phases (mid-log, late-log and stationary phase) with the highest average response to OMVs isolated during stationary phase and lowest from mid-log phase OMVs (Figure 1A). Notably, although all size classes (<3 kDa, 3-30 kDa, 30-100 kDa) induced metamorphosis in larvae of *H. elegans* (Figure 1B), the highest abundance of vesicles was also observed during stationary phase (Figure 1C).

**Figure 1.**
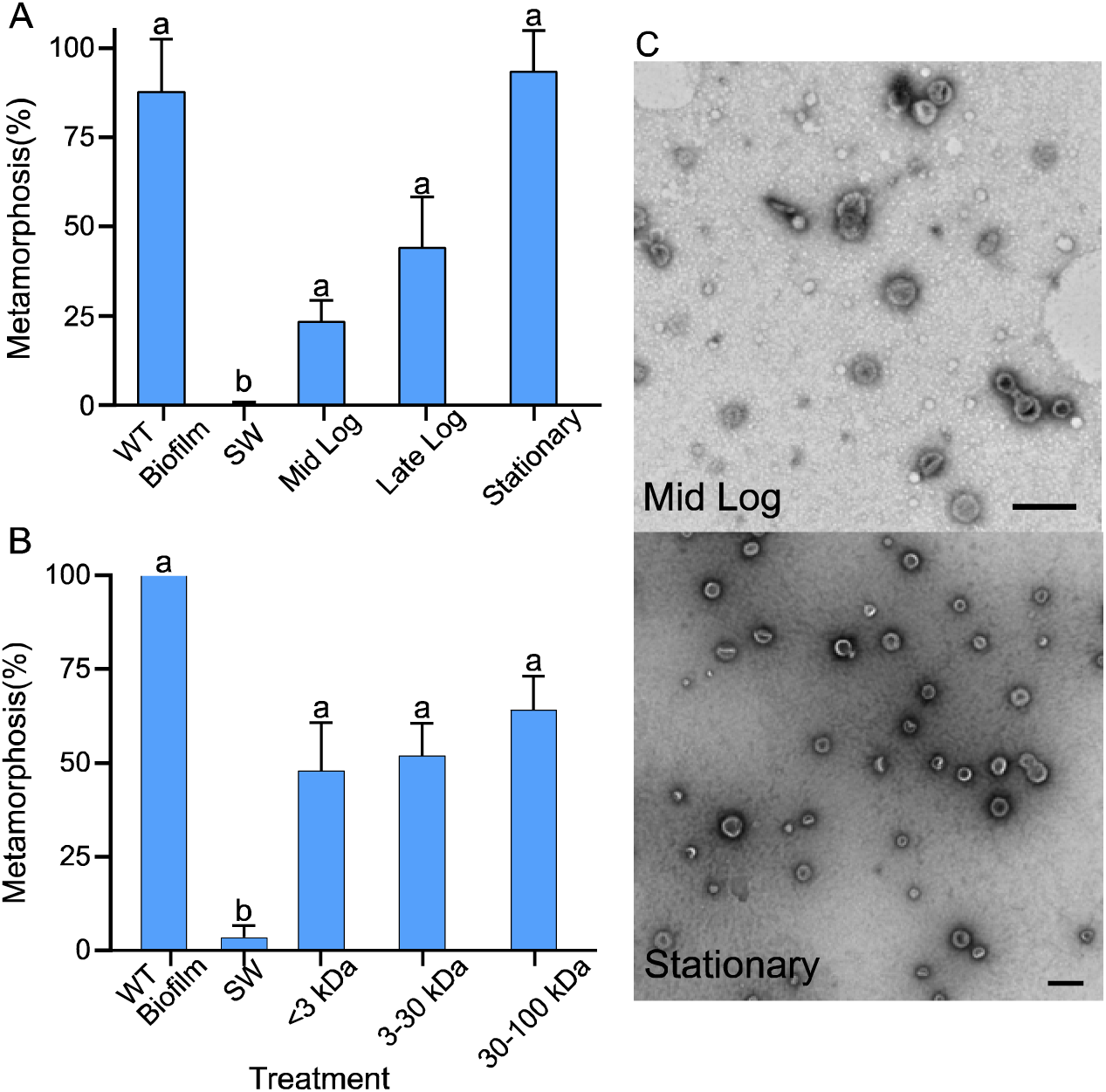
**Metamorphosis of larvae of *H. elegans* induced by outer membrane vesicles from *C. lytica.*** (A) Metamorphosis of larvae of *H. elegans* when exposed to OMVs from different size classes. (B) Metamorphosis of larvae of *H. elegans* when exposed to OMVs from each growth phase (mid log, late log and stationary). Metamorphosis was counted after 24 h exposure, with filtered seawater serving as a negative control, and a multispecies biofilm serving as a positive control. Letters show significant differences. (C) Negatively stained TEM image of OMVs from the mid log and stationary stage of a culture of *C. lytica*. Scale bar 200 nm.

### Enzymatic interrogations of OMVs for larval metamorphic cues: Nucleic acids and proteins

The enzymatic interrogations of OMVs from *C. lytica* with nucleases and proteases had no impact on the metamorphosis-inducing capacity of the OMVs (Figure 2A). The absence of nucleic acids in the OMV samples was confirmed by electrophoresis and UV absorbance, and the enzymatic destruction of proteins was confirmed by SDS-PAGE. Interestingly, both proteases had to be removed from the treatment by ultracentrifugation prior to larval assay, because the residual enzymes themselves induced metamorphosis (Fig S2).

**Figure 2.**
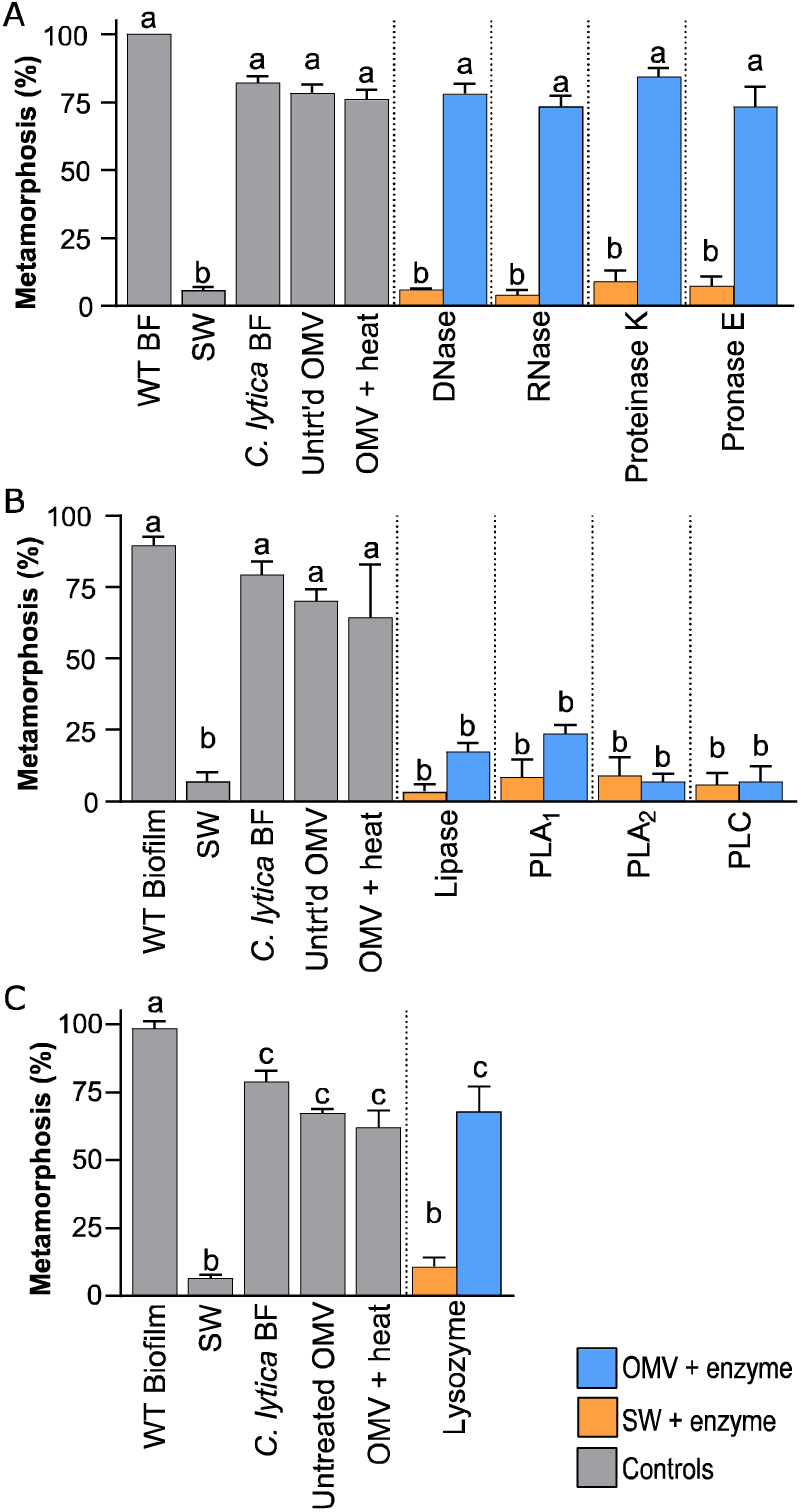
**Induction of metamorphosis in larvae of *H. elegans* from enzyme treated outer membrane vesicles (OMVs) from *C. lytica*.** OMVs and seawater (SW) controls were exposed to either (A) Nucleases (DNase (200 U) or RNase (200 U) and Proteases (trypsin (20 µg·ml^-1^) followed by either Proteinase K or Pronase E (100 µg·ml^-1^); (B) lipase (100 U), phospholipase A_1_ (PLA1; 100 U), phospholipase A_2_ (PLA2; 100 U) and phospholipase C (PLC; 100 U); (C) lysozyme (500 U). Metamorphosis was counted after 24 h exposure. Filtered seawater (SW) served as a negative control, and a multispecies biofilm, a *C. lytica* biofilm and an untreated *C. lytica* OMV sample were used as positive controls. Letters indicate significant differences.

### Enzymatic interrogations of OMVs for larval metamorphic cues: Lipids

The enzymatic interrogation of OMVs from *C. lytica* with lipases and phospholipases significantly decreased OMV induction of metamorphosis (Figure 2B). Metamorphosis after OMV treatment with Phospholipase A_1_, while still significantly higher than the treatments with all other lipases, was significantly lower than the untreated OMV and positive controls (p<0.05; Figure 2B). No molecular structure-activity relationship for the metamorphic cue can be predicted from the difference in reduction of metamorphosis between the phospholipase A_1_ and A_2_ treatments (Figure 2B). Phospholipase A_2_ is composed mostly of linear sheets and consequently will be subject to less steric hindrance than the barrel shaped phospholipase A_1_ (54). These results indicate the involvement of a bioactive lipid.

### Enzymatic interrogations of OMVs for larval metamorphic cues: Peptidoglycan

Enzymatic interrogation of OMVs from *C. lytica* with lysozyme (to break down peptidoglycan) had no impact on metamorphosis of larvae of *H. elegans* (Figure 2C).

### Bioassay guided fractionation of *C. lytica*

None of the HPLC fractions or purified compounds were as active as the parent samples. The compounds isolated through the bioassay-guided fractionation of *C. lytica* were all extracted from moderately active HPLC fractions of the cells of *C. lytica* (Figures S2-6). The amino acids (phenylalanine and leucine), as well as, niacin, thymidine, adenosine, and a 2,5-diketopiperazine of phenylalanine and proline were isolated and had minor levels of bioactivity (Figure 3). Bioactivity of isolated compounds was consistent with their commercial counterparts (Figures S5-6). It should be noted that induction of larval metamorphosis did not begin until after more than 6 h of exposure to any of the fractions, which is exceptionally slow.

**Figure 3.**
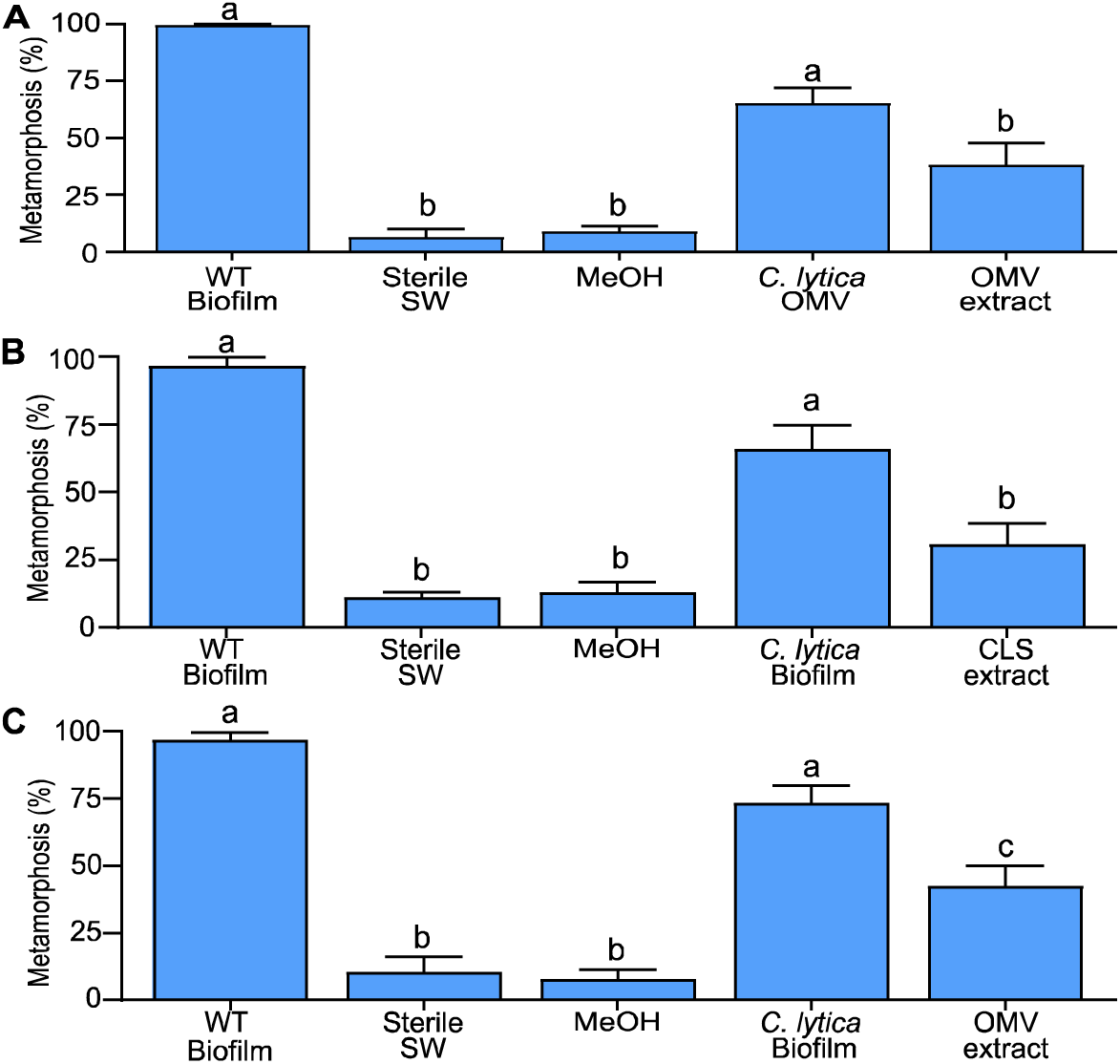
**Metamorphosis induction in larvae of *H. elegans* was consistently low from methanol extracts (10 µg·ml^-1^) of A) isolated OMV fraction; B) cell-free culture supernatant containing outer membrane vesicles (OMVs) of *C. lytica;* C) cell pellet.** Extracts were solubilized in ethanol, transferred to wells, evaporated to remove solvent and sterile seawater and larvae were added. Solvent control: ethanol, negative control: sterile seawater (SW), positive controls: Wild type (WT) biofilm and monospecific *C. lytica* biofilm.

Analysis of the genome of *C. lytica* with AntiSMASH (55, 56) identified one potential region of secondary metabolite significance from 445 027–465 863 bp. A protein -BLAST of this contig, identified it as a phytoene synthase (Table S1-2). Analysis of the genome of *C. lytica* with NaPDos (57) identified three potential ketosynthase domains, (Table S3), however, closer examination showed that these sequences are involved in fatty acid biosynthesis and are unrelated to the production of bioactive secondary metabolites.

### Analysis of cell envelope components for larval metamorphic cues

Isolated peptidoglycan at first appeared to induce settlement in larvae of *H. elegans*, but the activity disappeared upon treatment with lipase suggesting the isolated peptidoglycan was contaminated with a lipid. This activity was unaffected by treatment with lysozyme (Figure 4). Commercial muramyl dipeptides and muramyl tripeptides, tested across a concentration range of 1–100 µM, resulted in no metamorphosis of larvae of *H. elegans* when tested separately or in combination.

**Figure 4.**
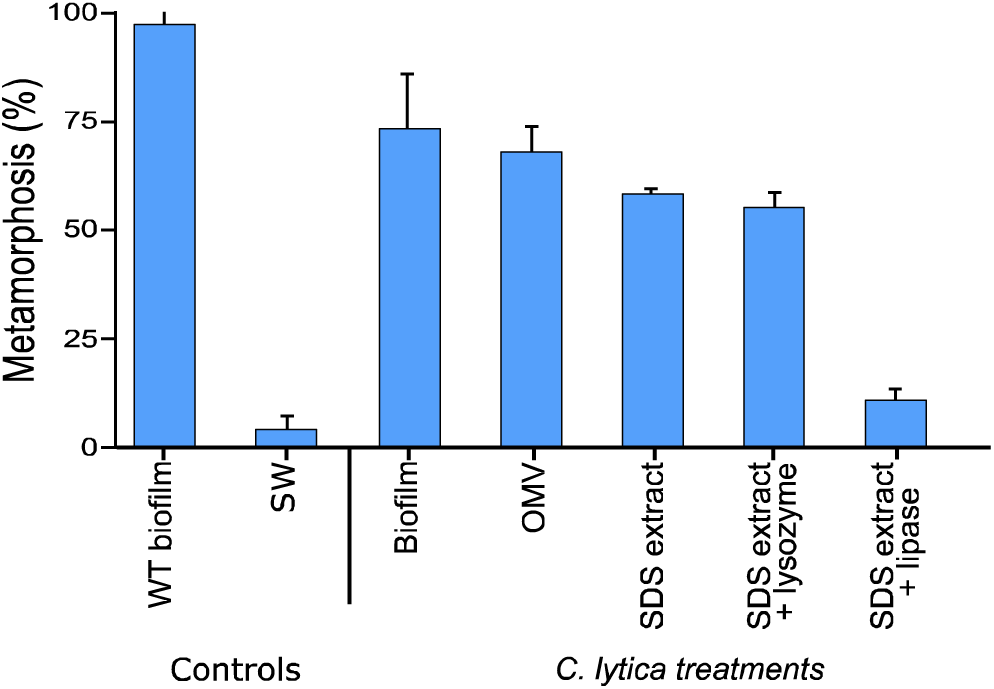
**A lipid that coextracted with peptidoglycan in the SDS extraction of cells of *C. lytica* induced metamorphosis in the larvae of *H. elegans.*** Metamorphosis was counted 24 h after exposure to the SDS extract (10 µg·ml^-1^), SDS extract that had been treated with lysozyme (500U) or SDS extract treated with lipase (100U). Negative control: sterile seawater (SW); positive controls a wild type (WT) biofilm, a *C. lytica* biofilm and *C. lytica* OMV sample.

Because lipases significantly reduced settlement induction by OMV samples and no significantly inductive lipophilic small molecules were found in the natural products isolations, our focus returned to the remaining major lipophilic component of the bacterial outer membrane, lipopolysaccharide.

Extracted and purified LPS successfully induced metamorphosis in the larvae of *H. elegans* (62%; Figure 5). Importantly, metamorphosis in larvae of H. elegans began less than one hour after exposure to LPS samples. A dose-response curve generated for this LPS fraction, yielded a peak of metamorphosis at 5 µg·ml^-1^. Metamorphosis was not induced when the concentration of the fraction dropped below 0.5 µg·ml^-1^, and toxicity and death were observed in larvae exposed to concentrations of LPS at or above 50 µg·ml^-1^ (Figure 5). Enzymatic treatment of the LPS sample was consistent with the results gained from the OMV experiments; only lipase reduced the induction of metamorphosis in larvae of *H*. elegans (Figure 5). The absence of peptidoglycan and proteins from this preparation was confirmed using SDS-PAGE, comparing Coomassie blue staining with silver staining (Figure 5). Additionally, the absence of proteins was indicated by a lack of absorption of the sample at 260 nm (supplementary information, part E). The absence of other contaminating glycolipids was confirmed with SDS-PAGE gels stained with the lipophilic stain Pro-Q Emerald 300 (Figure 5) as well as TLC developed with 5-hydroxy-1-tetralone. The Pro-Q Emerald 300 stain suggests that there could be at least 2 LPS molecules within the purified LPS sample.

**Figure 5.**
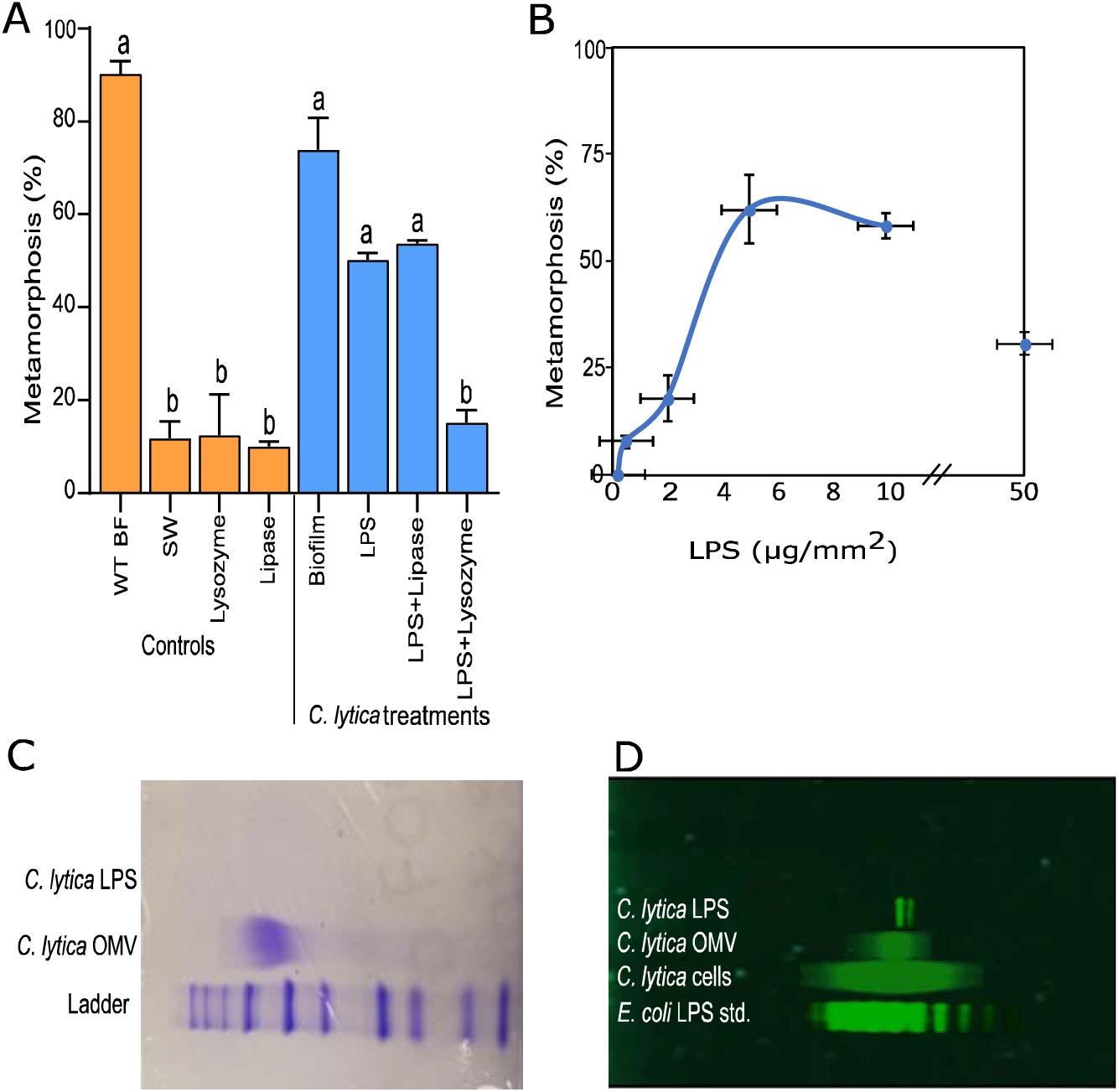
**LPS induces metamorphosis in larvae of *Hydroides elegans.*** (A) Induction of metamorphosis by LPS was not significantly affected by treatment with lysozyme or lipase (lysozyme (500 U), lipase (100 U); negative control: sterile filtered seawater (SW); positive control: wild-type (WT) biofilm and biofilm of *C. lytica*. Letters indicate significant differences). (B) Metamorphosis dose response curve of LPS isolated from *C. lytica*. (C) Coomassie Blue stained SDS-PAGE gel showing standard protein ladder, the presence of proteins in the OMVs isolated from *C. lytica*, and the absence of proteins in the extracted LPS sample. (D) SDS-PAGE gel stained with ProQ Emerald 300 showing *E. coli* LPS standard followed by boiled cells of *C. lytica*, OMVs from *C. lytica* and isolated LPS from *C. lytica*.

## Discussion

The broadly distributed marine polychaete *Hydroides elegans* serves as a useful proxy for undoubtedly thousands of benthic marine invertebrate species whose larvae settle and metamorphose in response to specific biofilm-dwelling bacteria (6, 12, 23, 24, 34, 58–61). However, it can be difficult to know whether bioactive chemicals identified in the laboratory are the same as those that act *in situ* or simply stimulate processes downstream of the receptor (62–66). The practice of isolating and identifying compounds involves the exposure of the larva to substances with which it may not naturally interact (62). While experimentally valuable, artificial induction of larval metamorphosis does not enable us to learn more of the true induction of metamorphosis that is so essential to benthic marine organisms. Fortunately, the wealth of knowledge accumulated for larval settlement conditions provides sound criteria for the acceptance or rejection of the ecological relevance of any isolated cue. Our conditions for accepting an ecologically relevant cue to induce metamorphosis in *H. elegans* require it to be bacterial in origin, fast acting and entrained to the biofilm (26). It was with this focus that we asked the question: what chemical component of marine biofilm bacterium *Cellulophaga lytica* cues the larva of *H. elegans* to settle and metamorphose?

OMVs, a key component for bacterial interactions with the environment, can be considered a secretion or a delivery system (42, 67). Previous studies have shown that OMVs produced during different culture growth stages or of different sizes can vary in composition and biological activity (42, 67, 68). Our data, however, demonstrated bioactivity in all growth stages, with a steady but non-significant increase over time, suggesting that the bioactive compound produced by *C. lytica* may be either constitutively produced or has a long half-life and may proportionately increase in cultures over time. To determine whether OMVs induce metamorphosis through the delivery of “cargo”, we subjected OMV fractions from *C. lytica* to a battery of enzymatic treatments and bioassay guided fractionation to identify components that impact larval settlement. Enzymatic interrogations of the OMVs strongly indicated that induction of metamorphosis was not due to a nucleic acid or protein cargo. Lipases, however, did inhibit induction of metamorphosis by the OMVs, triggering an investigation for the presence of a small bioactive lipid with bioassay-guided fractionation. Methanol and DCM extractions of OMV preparations followed by HPLC separation and bioassays failed to identify any bioactive lipids.

The bioassay-guided fractionation did identify a few low-activity common small molecules such as amino acids, but these were judged to not have ecological relevance. First, it is known that induction of metamorphosis can be stimulated in larvae of *H. elegans* by artificial inducers (25, 64). Previous studies into the induction of metamorphosis in *H. elegans* by dissolved free amino acids (DFAAs) was rejected due to delayed onset of metamorphosis, low levels of metamorphosis, non-specificity of DFAAs, undetectability of DFAAs in field studies and the presence of contaminating bacterial biofilms in the initial experimental set up (69–71). Second, the turbulent flow present at the field site, Pearl Harbor, would quickly dissipate FAAs and keep them well below inductive levels (29–31). Our results agree with these conclusions; when we tested amino acids, metamorphosis was rare and delayed. Additionally, all the extracts, HPLC fractions, and purified compounds were less active than their corresponding parent samples. For these reasons, we concluded that none of the low-activity metabolites from the bioassay-guided fractionations were involved in natural metamorphic induction. The *in-silico* searches further confirmed the results from bioassay-guided fractionation approach; no gene clusters or domains from known secondary metabolite biosynthetic pathways were identified that could transcribe compounds likely to induce complex larval metamorphosis.

The results of the enzymatic interrogations of the OMVs, in combination with the discovery that, in culture, OMVs were active settlement inducers regardless of their size or during which bacterial growth phase they were collected, suggested that OMVs induced metamorphosis because they were small pieces of cell membranes, rather than because they were acting as a delivery system. For this reason, we next focused our investigations on studies of the bacterial cell membranes. Cell membranes can contain a few larger lipid classes that may have been missed by the bioassay guided fractionation, including phospholipids, glycolipids, and glycosphingolipids.

Phospholipids, although considered a major source for activation-dependent signaling molecules (72), can be almost immediately dismissed from the list of potential settlement inducers. TLC chromatography of the active LPS sample showed a complete absence of phospholipid. Furthermore, not only is phospholipid signaling overwhelmingly intracellular (72, 73), but its activity also arises from secondary messenger molecules derived from the membrane phospholipids. These secondary messenger molecules are produced when phospholipases selectively hydrolyze phospholipids (72, 73). Consequently, the phospholipases should have increased production of the secondary messenger molecules and resulted in higher metamorphosis and not the significant reduction in metamorphosis that was observed (Fig. 2). Alternatively, the reduced activity of OMVs treated with phospholipase could reflect the requirement for an intact or negatively charged membrane (73). Selective externalization of phospholipids in the outer membrane can alter the biophysical properties of the membrane affecting their binding properties (72). Additionally, intact membranes may hold active cue molecules in particular conformations that would not occur if their components were in a free solution. If that conformation is required to activate the larval receptor, removal of the compound from the membrane would result in a similar loss of activity. This explanation, however, was quickly rejected by the discovery of inductive activity among the remaining classes of compounds obtained by extraction of the cell envelope. Rather, the inhibition of induction by lipase treatments is most likely the result of non-specific enzyme action.

The remaining classes of compounds included the important MAMP groups, peptidoglycan and lipopolysaccharide. Of these two groups, peptidoglycan has been shown to be an active component of non-pathogenic bacterial signaling in other systems (74, 75), and, for this reason, was chosen as the next component examined for its capacity to induce larval settlement. Results of larval assays utilizing isolated peptidoglycan, commercial peptidoglycan monomers or OMVs subjected to lysozyme treatments were all negative. These results are not entirely unexpected since the peptidoglycan layer resides beneath the outer membrane and would not be the first point of interaction between a larva and a bacterium or an OMV.

LPS, which could be described as a “prototypical MAMP”, was found to induce metamorphosis in larvae of *H. elegans* when provided at a concentration that reflects the concentration of cells in an active biofilm of *C. lytica* (24). In addition to being a major MAMP, LPS molecules comprise the vast outer portion of the outer membrane of Gram-negative bacteria, and, as such, would be the first bacterial molecules to interact with a larva upon contact, which is in line with how the process naturally occurs (26). Similarly, LPS forms the outer portion of the outer membrane of OMVs and is thus consistent with the observed activity of OMVs from *C. lytica*. It is important to note here that, although isolations of LPS can include residual traces of both peptidoglycan and other membrane lipids (phospholipids, glycosphingolipids etc.) (76), purity checks using both SDS-PAGE and TLC revealed none of these classes of compounds in the active LPS extract. Additionally, lipase and phospholipase treatments of the isolated LPS inhibited the inductive activity, consistent with the results from the OMV samples and suggesting that non-specific enzyme activity occurred. In other words, the discovery that the inductive cue for settlement and metamorphosis of larvae of *H. elegans* is comprised of bacterial LPS is consistent with all of the observations made previously and in this study.

While LPS may largely explain the widespread occurrence of inductive activity across Gram-negative biofilm bacteria, it cannot account for the same activity found in Gram-positive bacteria (34) which lack both an outer membrane and its LPS element. Instead, the cell envelope of Gram-positive bacteria consists of a thickened cell wall of peptidoglycan that can be populated by teichoic and lipoteichoic acids (LTA) (77, 78). LTA is a major MAMP of Gram-positive bacteria; it is a component of nearly all Gram-positive membranes, *as well as a functional equivalent to LPS* (79). Thus, we predict that LTA from the inductive Gram-positive bacteria will be the active moiety in induction of metamorphosis. However, given the great predominance of Gram-negative bacteria in marine biofilms (80) may render this possibility irrelevant, i.e., settlement induction in nature may always be by Gram-negative bacteria.

The strong immunogenicity of LPS, combined with its ubiquitous expression by Gram-negative bacteria and inherent structural variability, serve to make it both a likely target of evolutionarily conserved innate-immune receptors in larvae as well as suggest a possibly wide-spread mechanism for bacterial induction of marine-invertebrate metamorphosis. LPS is a highly variable macromolecule with this variability tied closely to bacterial taxonomic lines and growth conditions (81–85). Indeed, our purified LPS sample appears to contain at least 3 different LPS molecules, which is not unusual (86). LPS molecules are composed of three main components: lipid A (endotoxin); an inner core oligosaccharide (kDO); and an outer core oligosaccharide (O-antigen), with variation occurring primarily in the lipid A and O-antigen components (86). While beyond the scope of this paper, determining the location of the inductive portion of the macromolecule or whether it is a specific moiety of a specific LPS molecule produced by *C. lytica* will comprise our next focus of study. If variations in the structure of LPS are responsible for the observed inductive activity of different species and strains of bacteria, it can explain why many – but definitely not all – marine Gram-negative bacteria induce settlement and metamorphosis in the larvae of *H. elegans* (17, 23, 28) and other invertebrate species. It may also explain why not all strains of the same bacterium induce larvae of a single invertebrate species to settle (37). Most importantly, the inherent variation of LPS molecules could explain the differential settlement of different invertebrates in different habitats.

## Materials and Methods

### Larval culture and bioassays

Dioecious adult *H. elegans were* collected from docks in Pearl Harbor and maintained in the laboratory in continuously flowing, unfiltered seawater for several weeks without a noticeable decrease in fecundity. Gametes were obtained and larval cultured according to Nedved and Hadfield (23). Briefly, worms were induced to spawn by removal from their tubes. Gametes were mixed. Fertilization and development proceeded rapidly to a feeding trochophore stage in one day. Larvae were fed the single-celled alga *Isochrysis galbana* Tahitian Strain at a concentration of approximately 60,000 cells·ml^-1^. The larvae developed to the metamorphically competent nectochaete stage by day five and were utilized in metamorphosis experiments for 1-2 days. Because larvae of *H. elegans* will not metamorphose without exposure to bacterial biofilms or their products, wild, complex natural films developed on glass slides readily served as positive larval controls, and maintaining the larvae in double-filtered, autoclaved seawater (DFASW) served as negative larval controls. Assays also included a biofilm of *C. lytica* for reference. Assays were carried out in 24-well plates, and the percent of larvae that metamorphosed was determined at 20–24 hours. Significant differences (p<0.05) were calculated using Kruskal-Wallis followed by pairwise comparisons with false detection rate (FDR) correction (87).

### Bacterial culture and OMV isolation

*Cellulophaga lytica* were grown as previously described by Freckelton et al. (34) on ½ FSWt media (88). Briefly, cryogen bacterial stocks were streaked onto ½ FSWt agar plates and incubated for 24-48 h at 25°C. Single colonies were then used to inoculate 10 ml cultures of ½ FSWt broth and incubated for 12 h at 25°C. An aliquot of this starter culture was then used to inoculate 200 ml cultures and incubated for a further 14 h at 25°C. Cells were harvested by low-speed centrifugation (4000 *g*, 25 min 4°C). The cell-free broth was then filtered (0.2 µm, Millipore) to remove any remaining cells and then ultra-centrifuged (200,000 *g*, 2 h, 4°C) (34, 89) to isolate OMVs. All samples were examined by TEM using negatively stained preparations to confirm the presence of OMVs as previously described (34).

### Determination of relative concentration of OMVs

OMV production is highly dependent on growth conditions, and pelagic and biofilm bacterial populations can produce widely disparate amounts of OMVs (41, 42). The amount of OMVs produced by a bacterium can likewise be impacted by the growth phase of the population or the presence of other species. This variability results in difficulties in predicting an ecologically relevant concentration for testing. Consequently, isolated OMV suspensions were diluted and tested across three orders of magnitude. The lower (0.001X) dilution reflects that of the number of cells in 1 h biofilm (10^5^ cells/mm^2^) (24) and the highest concentration (1X) reflect the concentration used to inoculate those biofilms (10^8^ cells·ml^-1^). For clarity of presentations all figures present the data for the lowest dilution only, unless otherwise stated. Additional OMV dilutions are presented in the supplementary information part A. To maintain consistent relative concentration of OMVs between culture batches an aliquot of each sample was stained with the fluorescent lipophilic styryl dye FM-4-64 (Molecular Probes; T-3166) following the protocol described by Schooling and Beverage (46). Briefly, samples were suspended in Hank’s buffered saline solution (HBSS) and stained with 20 µM FM-4-64 in the dark for 20 min at 25°C. Fluorescence was read using a Molecular Devices SpectraMax M5 spectrophotometer (EX: 544 EM:680). Cholesterol was used to calibrate the dye’s response.

### Analysis of OMVs from different growth phases of cultures of *C. lytica*

Cultures of *C. lytica* were monitored for growth rate by measuring the OD_600_ of an aliquot hourly to determine the growth phase. This process was repeated with triplicate cultures, and samples were collected at early log, late log and stationary phases. OMVs were isolated from each sample time point as described above and diluted to reflect the concentration of a culture at 10^5^ cells·ml^-1^ (as above) in keeping with their FM-46-4 readings.

### Analysis of OMV size classes

Cell-free, filtered (0.2 µm), conditioned media was used to isolate OMVs. OMVs from *C. lytica* were separated into size classes by ultrafiltration (Millipore, Centricon Plus-70, 3 kDa, 30 kDa, 100 kDa). Aliquots (2 x 70 ml) of conditioned media were applied to each ultrafilter and then centrifuged (4 000 *g* x 30 min x 4°C). Aliquots of the retentate and filtrate were taken for metamorphosis assays and TEM analysis. The remaining filtrate was then applied to the next ultrafilter (100 kDa > 30 kDa > 3 kDa).

### Enzyme treatment of OMVs

Isolated OMVs were combined with enzymes (nucleases, proteases, lipases, or lysozyme) and incubated in a water bath at the optimal temperatures and time periods described in Table 1. Lysozyme is an enzyme that catalyzes the breakdown of 1,4-beta-linkages between N-acetylmuramic acid and N-acetyl-D-glucosamine residues in peptidoglycan (90). Enzyme action was halted by heat treatment alone, except for the proteases which had to be removed from the treatment by ultracentrifugation prior to larval assay, because the residual enzymes themselves induced metamorphosis (supplementary information part B, Fig S1). For each enzyme or set of enzymes, a seawater control was also performed to ensure that the responses observed were not a result of the enzymes or heat treatments alone. Successes of nuclease treatments were assessed by spectrometry and electrophoresis. Successes of protease treatments were assessed by spectrometry and SDS-PAGE.

**Table 1:**
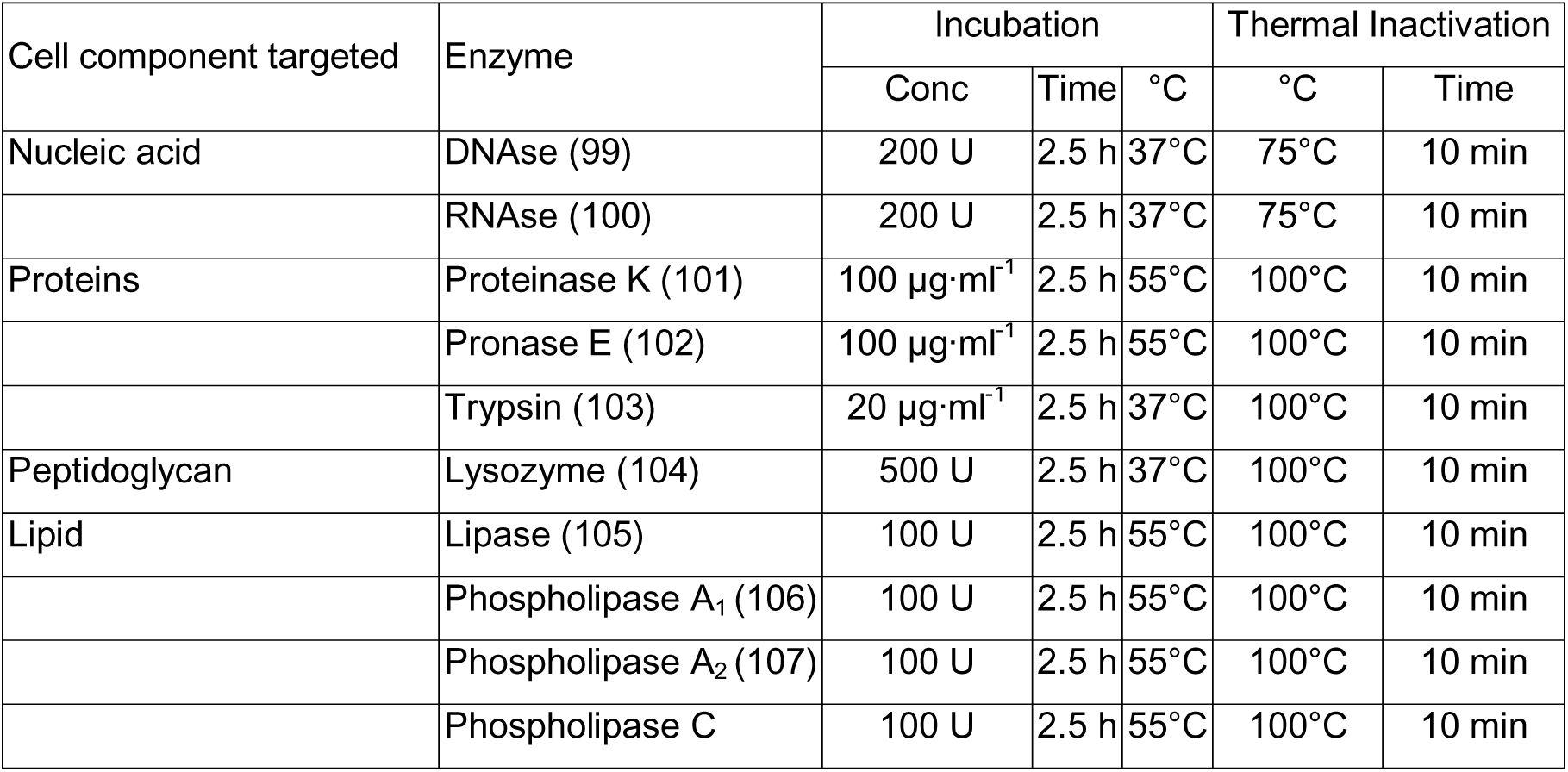
Enzyme treatments of OMVs from *C. lytica*.

| Cell component targeted | Enzyme | Incubation |  |  | Thermal Inactivation |  |
| --- | --- | --- | --- | --- | --- | --- |
|  |  | Conc | Time | °C | °C | Time |
| Nucleic acid | DNAse (99) | 200 U | 2.5 h | 37°C | 75°C | 10 min |
|  | RNAse (100) | 200 U | 2.5 h | 37°C | 75°C | 10 min |
| Proteins | Proteinase K (101) | 100 µg·ml<sup>-1</sup> | 2.5 h | 55°C | 100°C | 10 min |
|  | Pronase E (102) | 100 µg·ml<sup>-1</sup> | 2.5 h | 55°C | 100°C | 10 min |
|  | Trypsin (103) | 20 µg·ml<sup>-1</sup> | 2.5 h | 37°C | 100°C | 10 min |
| Peptidoglycan | Lysozyme (104) | 500 U | 2.5 h | 37°C | 100°C | 10 min |
| Lipid | Lipase (105) | 100 U | 2.5 h | 55°C | 100°C | 10 min |
|  | Phospholipase A<sub>1</sub> (106) | 100 U | 2.5 h | 55°C | 100°C | 10 min |
|  | Phospholipase A<sub>2</sub> (107) | 100 U | 2.5 h | 55°C | 100°C | 10 min |
|  | Phospholipase C | 100 U | 2.5 h | 55°C | 100°C | 10 min |

### Bioassay-guided fractionation of *C. lytica*

Cell pellets (CLP), cell-free supernates (CLS) and OMVs were extracted with organic solvents directly to identify metamorphosis-inducing molecules. CLP and OMV samples were extracted with methanol (MeOH) followed by a dichloromethane-methanol mixture (DCM: MeOH, 1:1). Methanol was used as the first extracting solvent to both quickly quench the cells and to remove any residual water. As this aqueous methanol extract would have primarily targeted polar molecules, this extraction was followed by a DCM: MeOH to recover any non-polar molecules. Some non-polar molecules will be extracted by both solvent systems but should exist in higher concentration in the DCM: MeOH extract. CLS (3.5 L) was extracted by liquid: liquid partition with ethyl acetate (EtOAc), with nonpolar compounds extracted into the EtOAc layer.

The initial OMV extracts were subjected to metamorphosis bioassays first at 10 µg·ml^-1^. Any samples that produced a toxic response in the larvae were re-assayed at 1 µg·ml^-1^ and 0.1 µg·ml^-1^. Extracts were added to wells by dissolving in MeOH at the concentration required for the addition of no more than 50 µL of MeOH to sample wells. MeOH was allowed to evaporate before drying overnight in a lyophilizer to remove any residual MeOH. A MeOH solvent control was added to the assay to ensure any observed induction of metamorphosis was not due to the residual presence of MeOH.

Additionally, each extract (CLP, CLS, OMV) was subjected to reverse phase HPLC (Thermo Scientific 3000), and all fractions were subjected to metamorphosis bioassay with larvae of *H. elegans*. CLP HPLC fractions were generated for bioactivity testing through an acetonitrile (ACN) gradient (10-25%, 20 mins; 25-40%, 10 mins; 40-100%, 20 mins; 100%, 10 mins). Active fractions were further purified using HPLC (Phenomenex, Luna, 5 µm, C18, 100 Å, 250 x 100 mm, MeOH 10%, 30 mins). For CLS, extracts were fractionated using a phenyl-hexyl column (Phenomenex, Luna, 5 µm, Phenyl-Hexyl, 250 x 100 mm) and a water: acetonitrile gradient. ^1^H NMR and LC-ESI-MS data were collected on all active fractions using a Bruker (400 MHz) and an Agilent 1260 HPLC system coupled to a 6120 quadrupole LC-MS spectrometer, respectively. Active fractions underwent ^1^H NMR and HR-MS analysis to assess purity, and, if fractions were still mixtures of compounds, they underwent additional fractionation. This process was repeated until pure fractions were obtained. Specific purification details of isolated compounds are provided in the supplementary information part C.

### *In silico* exploration for the presence of secondary metabolite gene clusters in the genome of *C. lytica*

AntiSMASH and NaPDos are bioinformatic platforms that allow genome mining for secondary metabolite biosynthesis gene clusters in bacterial genomes (55, 56) and allow rapid genome-wide identification of secondary metabolite genes, C- or KS- domains or amino acids. *In silico* analyses of the genome of *C. lytica* (CP009239.1) using both AntiSMASH 5.0 (56, 57) and NaPDos (57) were run with default, relaxed parameters to determine the potential for *C. lytica* to produce secondary metabolites (supplementary information part D, Tables S3-5).

### Extraction of cell-envelope peptidoglycan of *C. lytica*

Peptidoglycans were obtained from cultures of *C. lytica* with a protocol adapted from Leulier et al. (76). Culture growth was halted by chilling cultures on ice to 4°C. Cells were collected by centrifugation, washed once with a cold 3.5% NaCl solution and centrifuged again. Cells were then suspended in hot (95°C) aqueous SDS solution (4%) for 30 min and then cooled to room temperature (1 h). The SDS was removed by ultracentrifugation (20000 *xg;* 40 mins) and the pellet was resuspended in water and washed again for a total of 5 washes. The pellet was then resuspended in water and incubated with both DNase I, RNase, Proteinase K, Pronase E and Trypsin (Table 1) to remove any contaminating proteins or nucleic acids. Enzymes were removed following the same ultracentrifuge protocol described above. Metamorphic activity in the peptidoglycan extract was further interrogated with either a lysozyme or lipase treatment. Induction of metamorphosis in larvae of *H. elegans was conducted* before and after lipase and lysozyme treatments.

### Bioactivity testing of commercially available cell-envelope components

Peptidoglycan monomers in the form of muramyl dipeptides and muramyl tripeptides (Sigma Aldrich) were tested for induction of metamorphosis in larvae of *H. elegans*, individually and across a range of concentrations (0.1–10 µM), by dissolving them in water. Additionally, a solution containing 10 µM concentrations of both compounds was also screened for metamorphic induction. LPS from *Escherichia coli* 055:B5*, Salmonella enterica* serotype enteritidis and *Pseudomonas aeruginosa* 10 (Sigma Aldrich) were tested for induction of metamorphosis in larvae of *H. elegans*, individually across a range of concentrations (0.1–10 µM), by dissolving them in DMSO adding them to the sample wells before freezing and lyophilizing overnight to remove any residual solvent. A solvent control was included in the assay to ensure any observed induction of metamorphosis was not due to the presence of residual solvent. All commercial LPSs were checked for contaminants with thin layer chromatography and ultraviolet spectroscopy.

### Extraction of cell-envelope lipopolysaccharide of *C. lytica*

Lipopolysaccharides (LPS) were extracted using the Westphal hot phenol method (91) as updated by Apicella et al. (92, 93). The LPS fraction was assessed for metamorphic induction in larvae of *H. elegans* across a range of concentrations determined from the inoculation density of metamorphically inductive biofilms of *C. lytica* (10^5^ cells·mm^-2^) and estimates of the amount of LPS present per cell range from 25-62 fg·cell^-1^ (94–96). This calculation estimates that active biofilms of *C. lytica* contain between 2 and 6 µg LPS per well in a metamorphosis assay. Consequently, a range from 0.5 µg·ml^-1^ to 10 µg·ml^-1^ LPS was utilized to develop a dose-response series for assessment of induction of metamorphosis in larvae of *H. elegans*.

To reduce issues with solubility of the LPS extract and to determine a weight-for-volume concentration, an aliquot of the concentrated LPS sample was dried completely using a speed-vac. This mass was then used to calculate the concentration of the remaining solution. The LPS sample was also subjected to enzymatic interrogation with lipase, lysozyme and pronase E as described above.

### Analysis of LPS extract purity

Aqueous LPS solutions were checked for contaminating proteins by measuring the absorption at 260 nm. Electrophoretic behavior and additional protein contamination checks were performed by SDS-PAGE (Coomassie Blue and Silver staining) (76, 97). The presence of glycolipids was identified with additional SDS-PAGE gels stained with Invitrogen Pro-Q™ Emerald 300 Lipopolysaccharide Gel Stain Kit according to manufacturer’s instructions. The presence of contaminating lipid classes in the extracted LPS was checked by utilizing the rapid fluorometric micro-detection system of Watanabe and Mizuta (98). Briefly, the LPS extract was run on an alumina backed silica TLC plate (AAdvance Instruments, 60Å F254) alongside LPS and glucose standards, as well as the phospholipid standard, phosphatidyl ethanolamine (Sigma Aldritch). The TLC was developed by heating with acidified 5-hydroxy-1-tetralone (98). Under UV light (365 nm) hexoses, including oligosaccharides and polysaccharides containing hexoses appear as green spots; acidic and neutral glycosphingolipids appear as a yellow spot and phospholipids appear as blue spots.

### Examination of bacterial preparations with transmission electron microscopy (TEM)

Bacterial preparations and cell-free preparations were applied to glow-discharged formvar grids, negatively stained with 2% uranyl acetate aqueous solution and air dried. Samples were imaged using a 120 kV Hitachi HT7700 electron microscope with an AMT XR-41 camera.

## Acknowledgments

This research was funded in part by the Gordon and Betty Moore Foundation through Grant GBMF5009 and the Office of Naval Research grant no. N00014-08-1-2658 to MGH. The authors especially thank Prof. Michael Apicella for his invaluable advice regarding all things pertaining to LPS and Mrs. Tina M. Carvalho at the Biological Electron Microscopy Facility, University of Hawai□i at Mānoa for her technical advice. Previous lab members, Nidhi Vijayan, Audrey Asahina and Shaun Hennings assisted us in many ways, which we greatly appreciate.

## Supplementary Information Text

### Part A: Growth phase analysis of OMVs

**Figure. S1.**
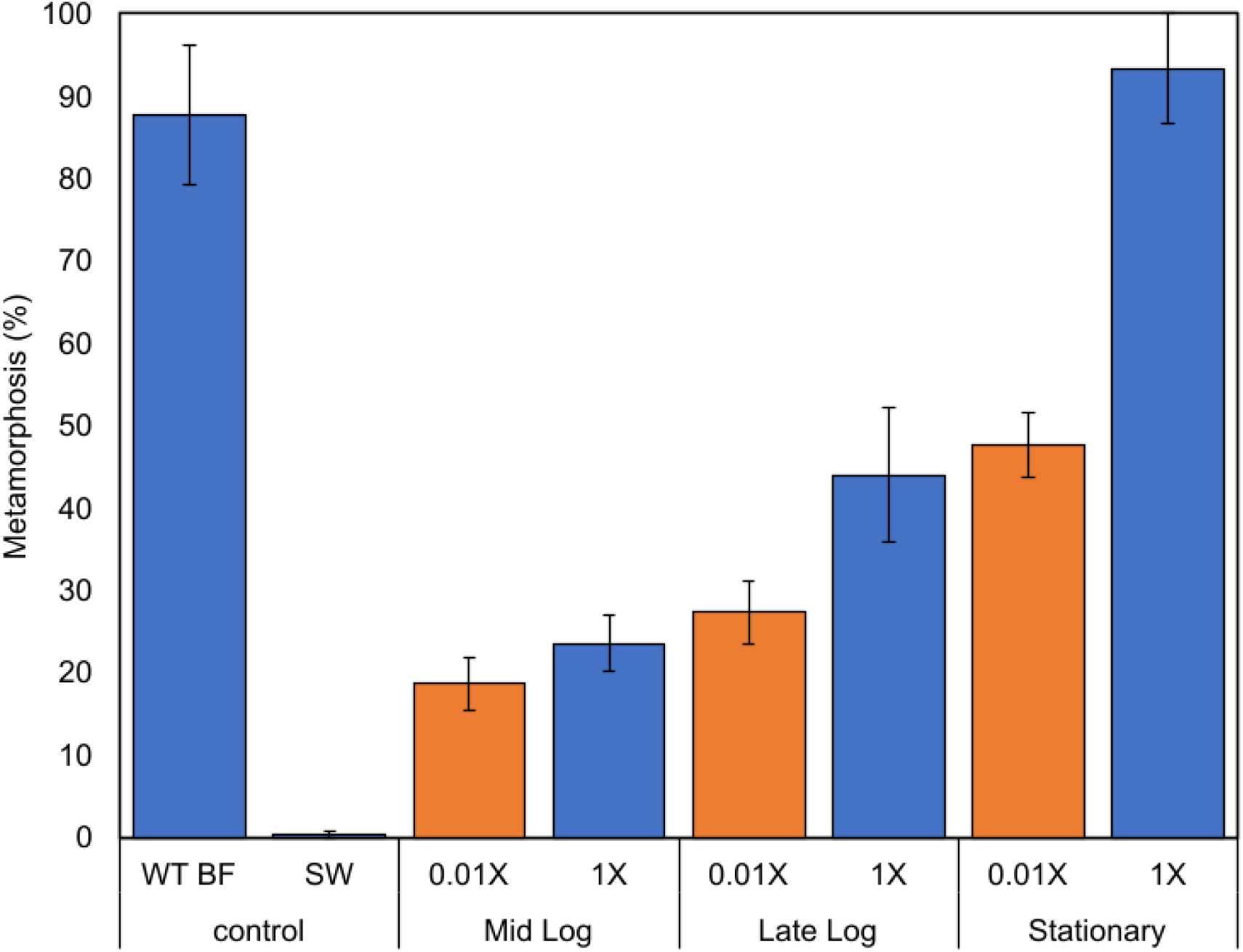
**Metamorphosis of larvae of *H. elegans* induced by outer membrane vesicles from a culture of *C. lytica* at different growth phases.** Metamorphosis of larvae of *H. elegans* when exposed to OMVs from each growth phase (mid log, late log and stationary). 1X is equivalent to the concentration of OMVs found in an inductive 10^8^ cells·ml^-1^ biofilm. 0.01X is a 1/100^th^ dilution. Metamorphosis was counted after 24 h exposure, with filtered seawater serving as a negative control, and a multispecies biofilm serving as a positive control. Letters show significant differences.

### Part B. Enzyme treatment of OMVs

**Figure. S2.**
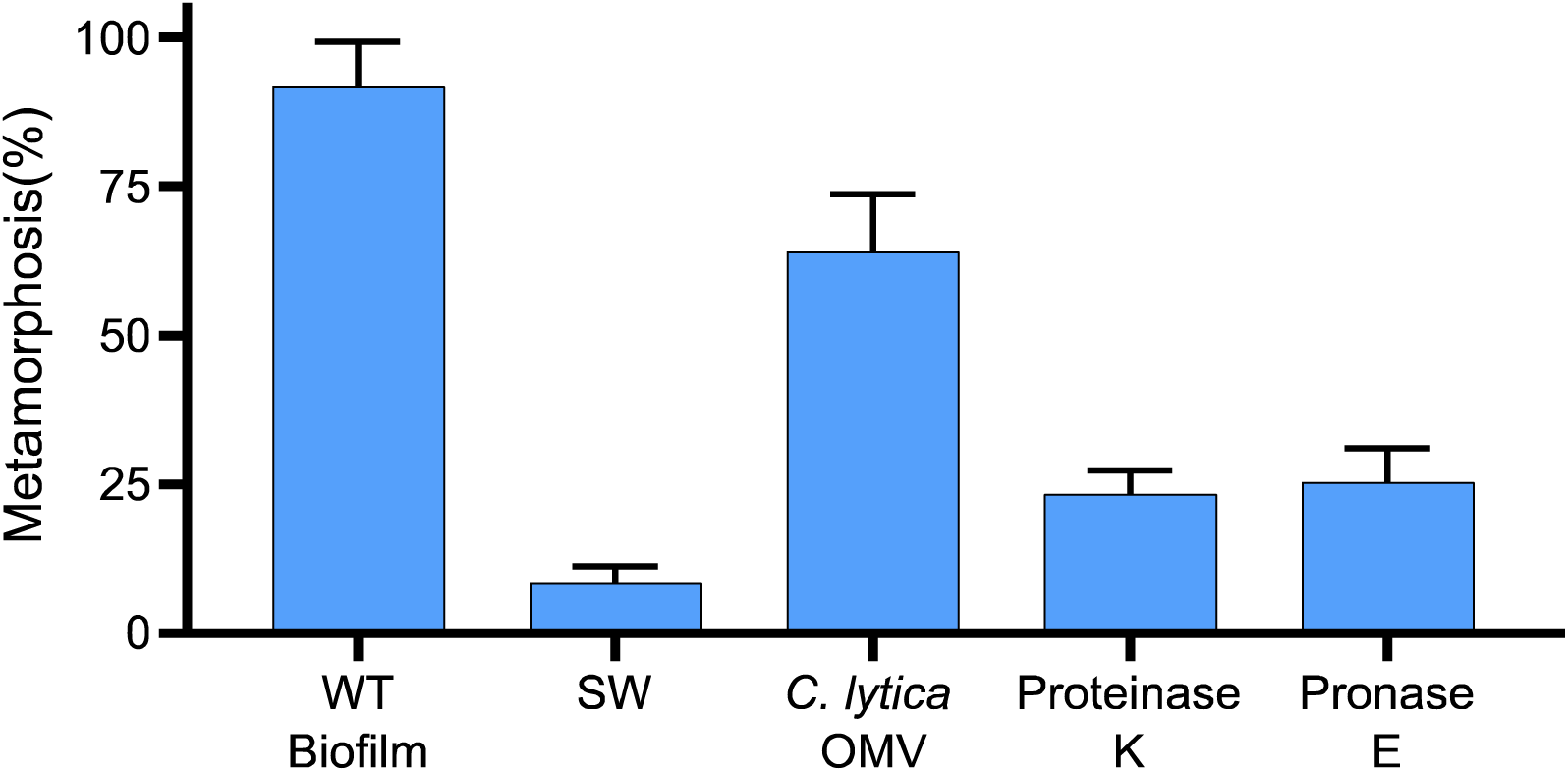
**Proteases induce metamorphosis in larvae of *C. lytica*. Larvae of *H. elegans* were exposed to either Proteinase K (5 U) or Pronase E (5 U).** Metamorphosis was counted after 24 h exposure. Filtered seawater (SW) served as a negative control, and a multispecies biofilm and untreated OMVs were used as positive controls.

### Part C: Bioassay-guided fractionation of *C. lytica*

#### 1. Small molecule analysis of cell pellets from *C. lytica*

Cell pellets of *C. lytica* were extracted with MeOH twice followed by repeated extraction with MeOH: DCM (1:1). The MeOH: DCM extract was subjected to semi-preparative reverse chromatography HPLC (C18, ACN: 10-25%, 20 mins; 25-40%, 10 mins; 40-100%, 20 mins; 100% 10 mins). Fractions were collected every minute, dried and resuspended at 20 mg/ml filtered seawater and tested for settlement-induction activity at a concentration of 10 µg/ml. Activity was detected in fraction 7 and fraction 29; fraction CLP-29 did not contain enough material or any indicative absorbance to pursue further.

Semi-preparative reverse chromatography of CLP7 (C18; MeOH 10%, 30 mins) yielded activity in the 2^nd^ and 4^th^ fractions (Fig S3A). LC-Mass spectrometry and ^1^H NMR spectroscopy were consistent with the presence of phenylalanine and adenosine (Fig S3B-D). To confirm the activity and further elucidate the stereochemistry, both isolated and commercial versions of adenosine (Sigma-Aldritch) and phenylalanine (D and L; Sigma-Aldritch) were tested for metamorphic induction (Fig S4). Additional commercially available compounds (thymidine, leucine (D and L) and niacin) were also tested (Fig S5).

#### 2. Small molecule analysis of cell-free supernatant from *C. lytica* containing OMVs

Conditioned media of *C. lytica* cultures (3.7 L) were extracted with ethyl acetate (EtOAc) and dried (363 mg). The EtOAc extract was subjected to semi-preparative reverse chromatography HPLC (C18, ACN: 10-25%, 20 mins; 25-40%, 10 mins; 40-100%, 20 mins; 100% 10 mins). Fractions were collected every minute, dried and resuspended at 20 mg/ml ethanol and tested for metamorphic activity at a concentration of (10 µg/ml). An additional solvent (ethanol) control was included in bioassay. Activity was identified in the 25^th^ and 26^th^ fractions which were combined. LC-MS revealed a compound consistent with the presence of a diketopiperazine of phenylalanine and proline (Cyclo(- Phe-Pro)). Isolated and commercial versions of this compound were subjected to metamorphosis assay as previously described (Fig S6).

**Figure. S3.**
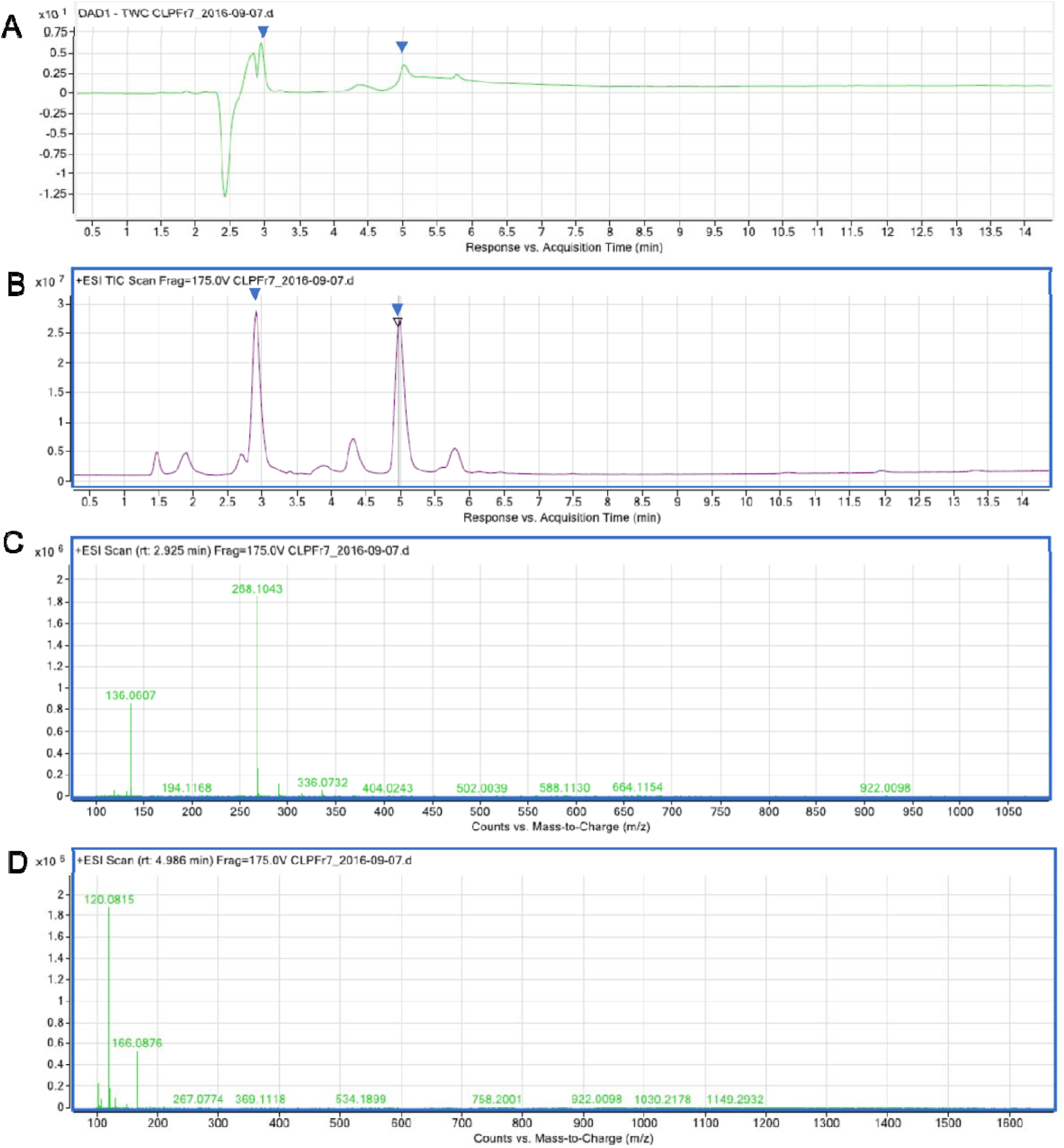
Spectral confirmation of isolation of adenosine and phenylalanine from most active cell pellet fraction (CLP7). (A) UV-Vis absorbance chromatogram B) Total Ion Chromatogram (TIC) positive mode, Ions were detected within a mass range m/z 100-1100. (C) Extracted ion chromatogram of the active peak at 2.9 min yielded m/z of 268.1043.(D) Extracted ion chromatogram of the active peak at 4.9 min yielded m/z of 123.0459.

**Figure. S4.**
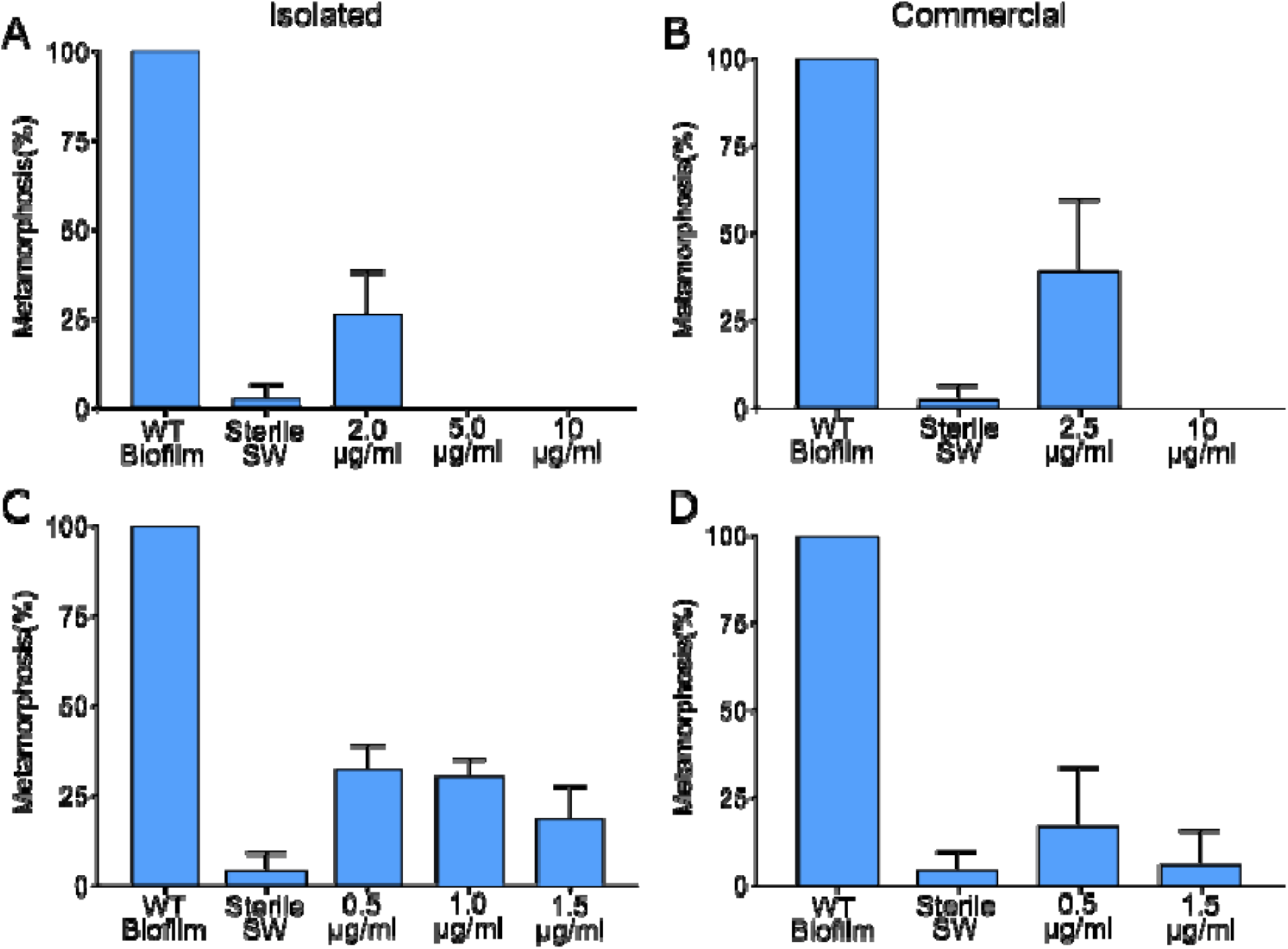
Metamorphosis of larvae of *H. elegans* when exposed to isolated (A) and commercial (C) adenosine and isolated (B) and commercial (D) phenylalanine from Fraction 7 of the cell pellet for 24 hrs. Positive Control: Wild Type (WT) Biofilm. Negative Control: Sterile Seawater (SW).

**Figure. S5.**
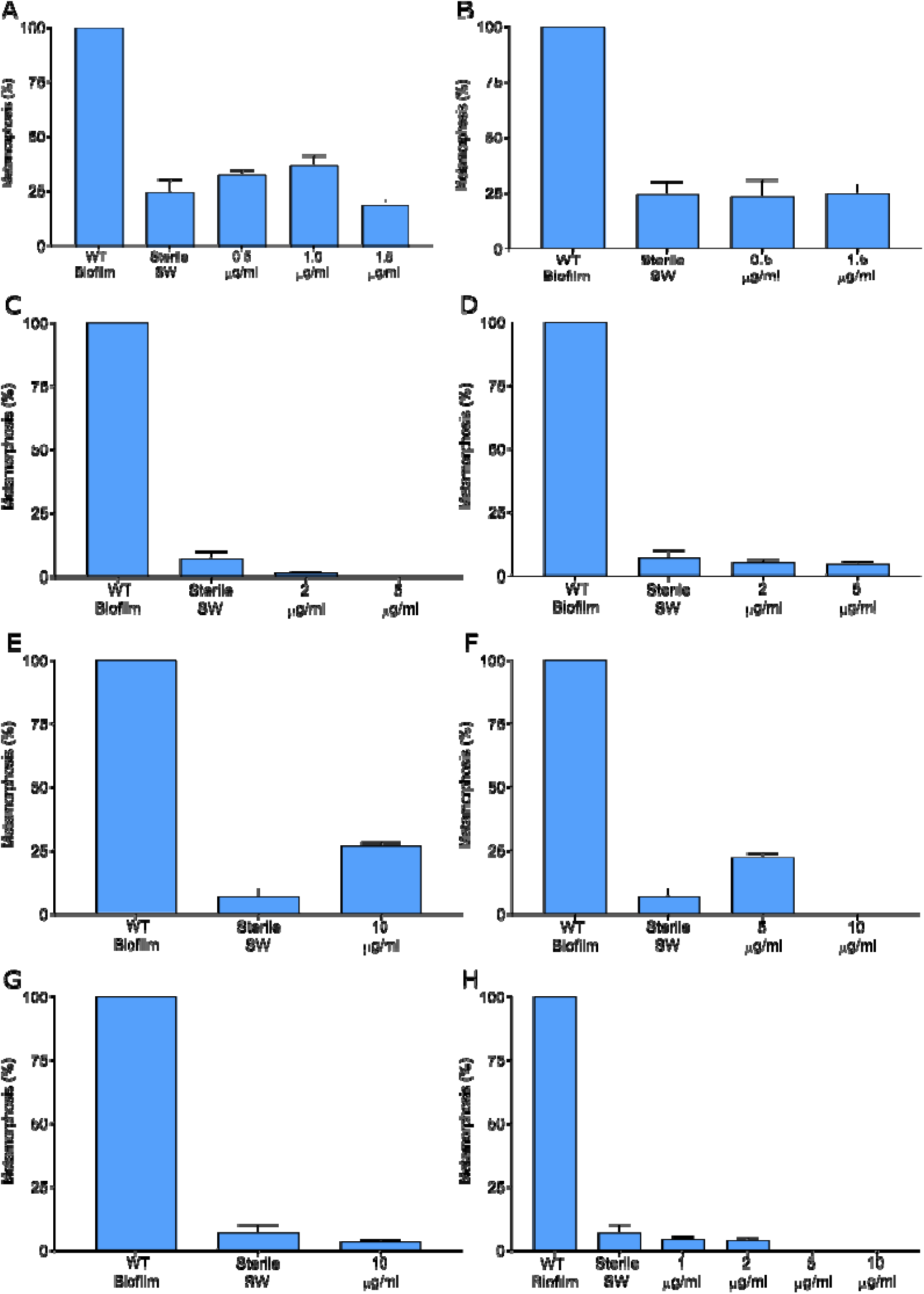
Metamorphosis of larvae of *H. elegans* when exposed to commercial versions of compounds detected in fraction 7 for 24 hrs. A) Adenosine; B) DL-Phenylalanine; C) L-Phenylalanine; D) D-Phenylalanine; E) Niacin; F) Thymidine; G) L=Leucine; H) D-Leucine. Positive Control: Wild Type (WT) Biofilm. Negative Control: Sterile Seawater (SW).

**Figure. S6.**
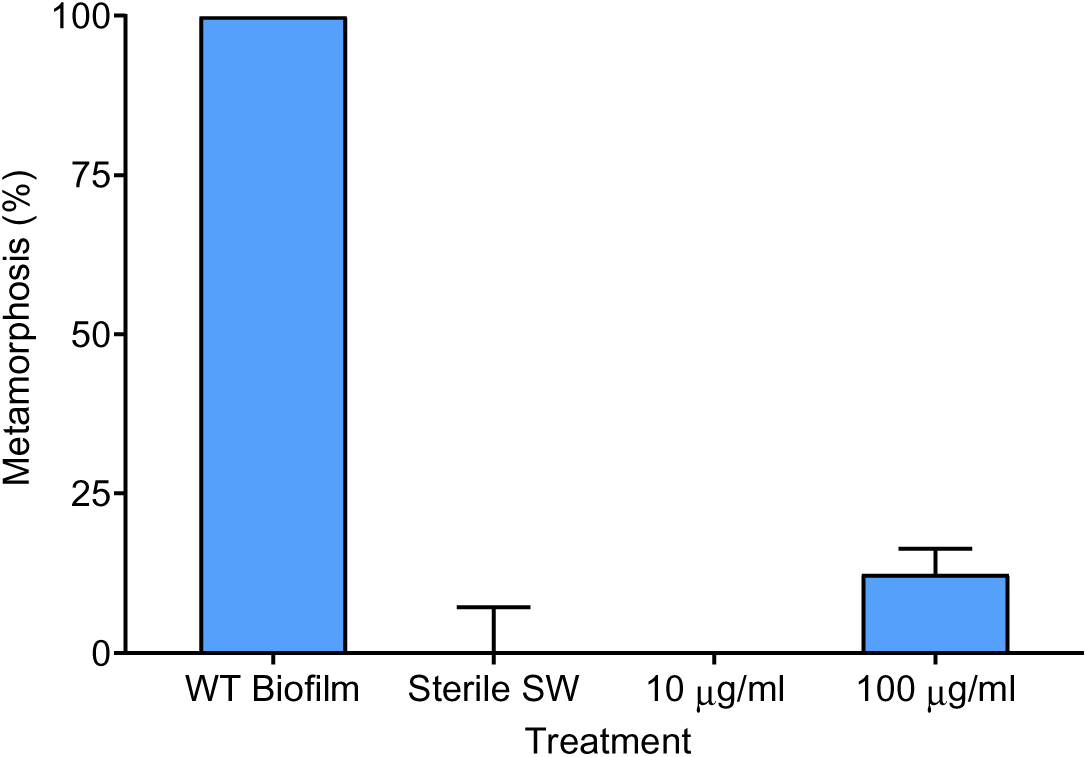
Metamorphosis of larvae of *H. elegans* when exposed to commercial version of the diketopiperizine of phenylalanine and proline (Cyclo(-Phe-Pro) detected in fraction 25 of the supernatant from a *C. lytica* culture for 24 hrs. Positive Control: Wild Type (WT) Biofilm. Negative Control: Sterile Seawater (SW).

### Part D. Genome Mining for Secondary Metabolites

Analysis of the genome of *C. lytica* for the presence of secondary metabolite biosynthesis gene clusters revealed a single contig *(ctg1_392)* located between 455,027 - 455,863. A protein blast of this contig, identified it as a phytoene synthase (Table S1). The Minimum Information about a Biosynthetic Gene cluster (MIBiG) failed to match any enzymes with greater than 50% sequence identity (Table S2). Similar low matches were generated using NaPDos (Table S3).

**Table S1.**
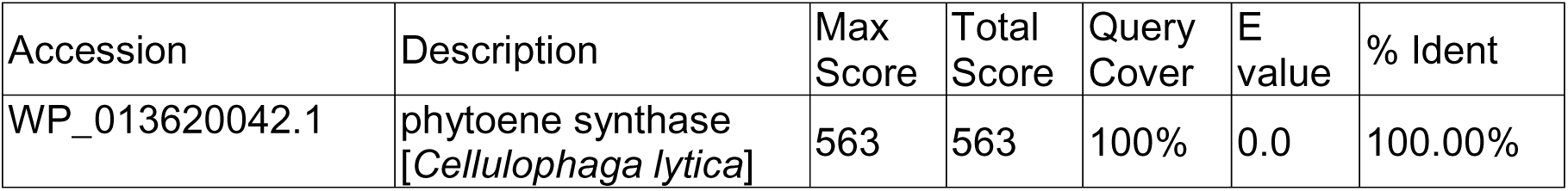
NCBI BlastP on ctg1_392 from genome of *C. lytica* generated by AntiSMASH (1)

**Table S2.**
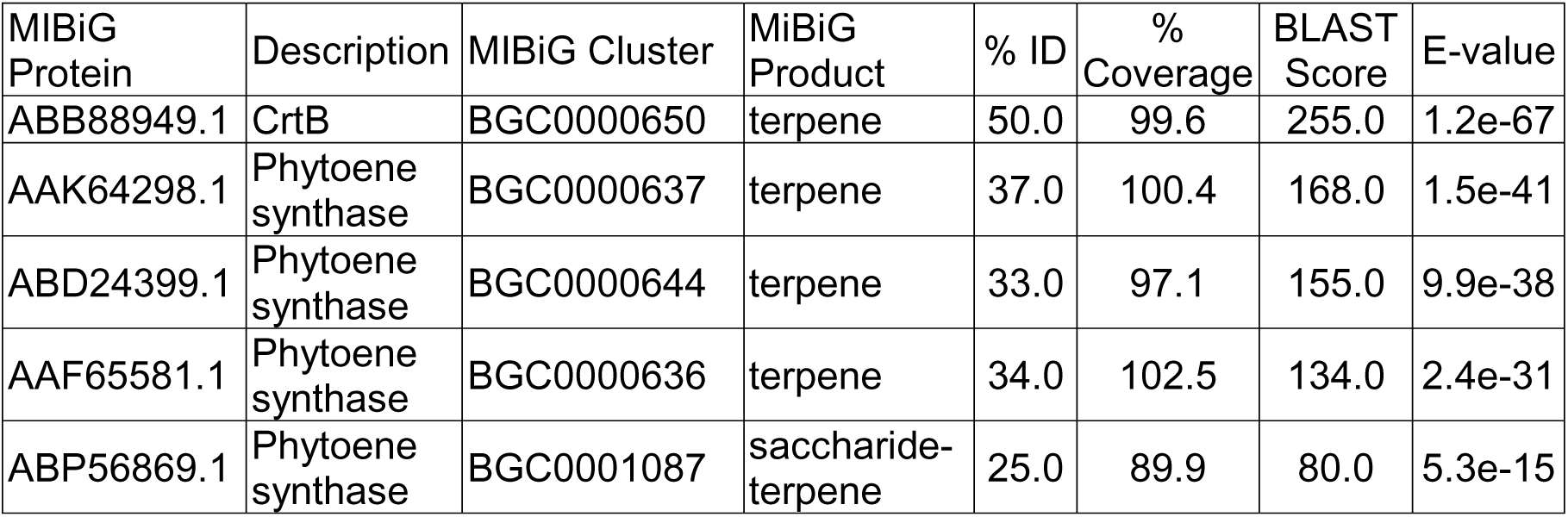
Minimum Information about a Biosynthetic Gene cluster (MIBiG) search results from genome of *C. lytica* generated by AntiSMASH (1)

**Table S3:**
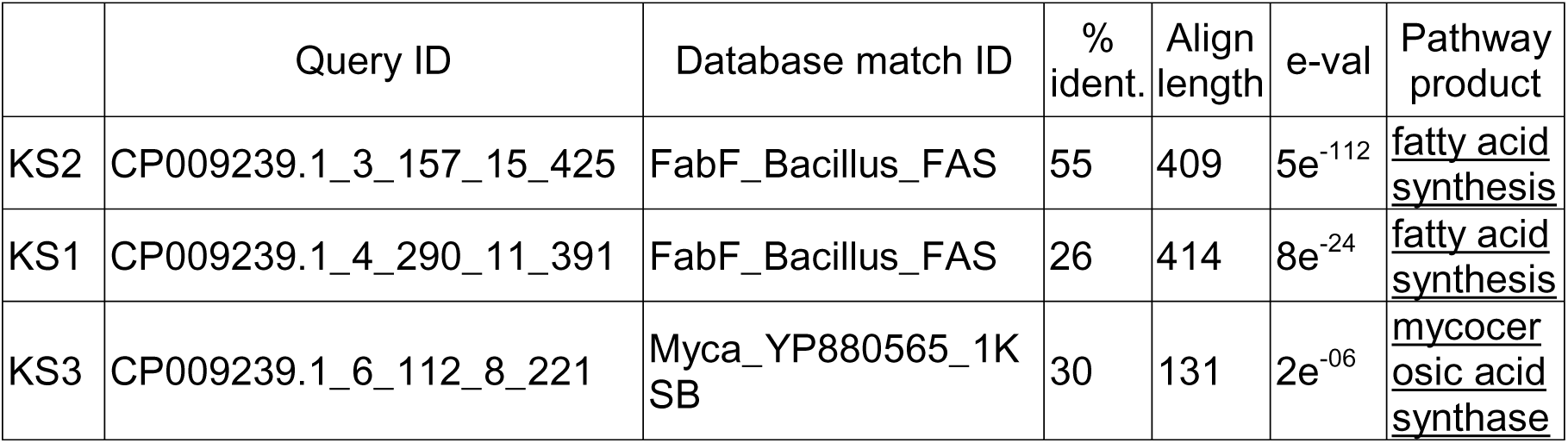
Polyketide Synthase (KS) domains within the genome of *C. lytica* generated by NaPDos (2). Search was run with relaxed parameters.

### Part E: LPS purity checks

**Figure S7.**
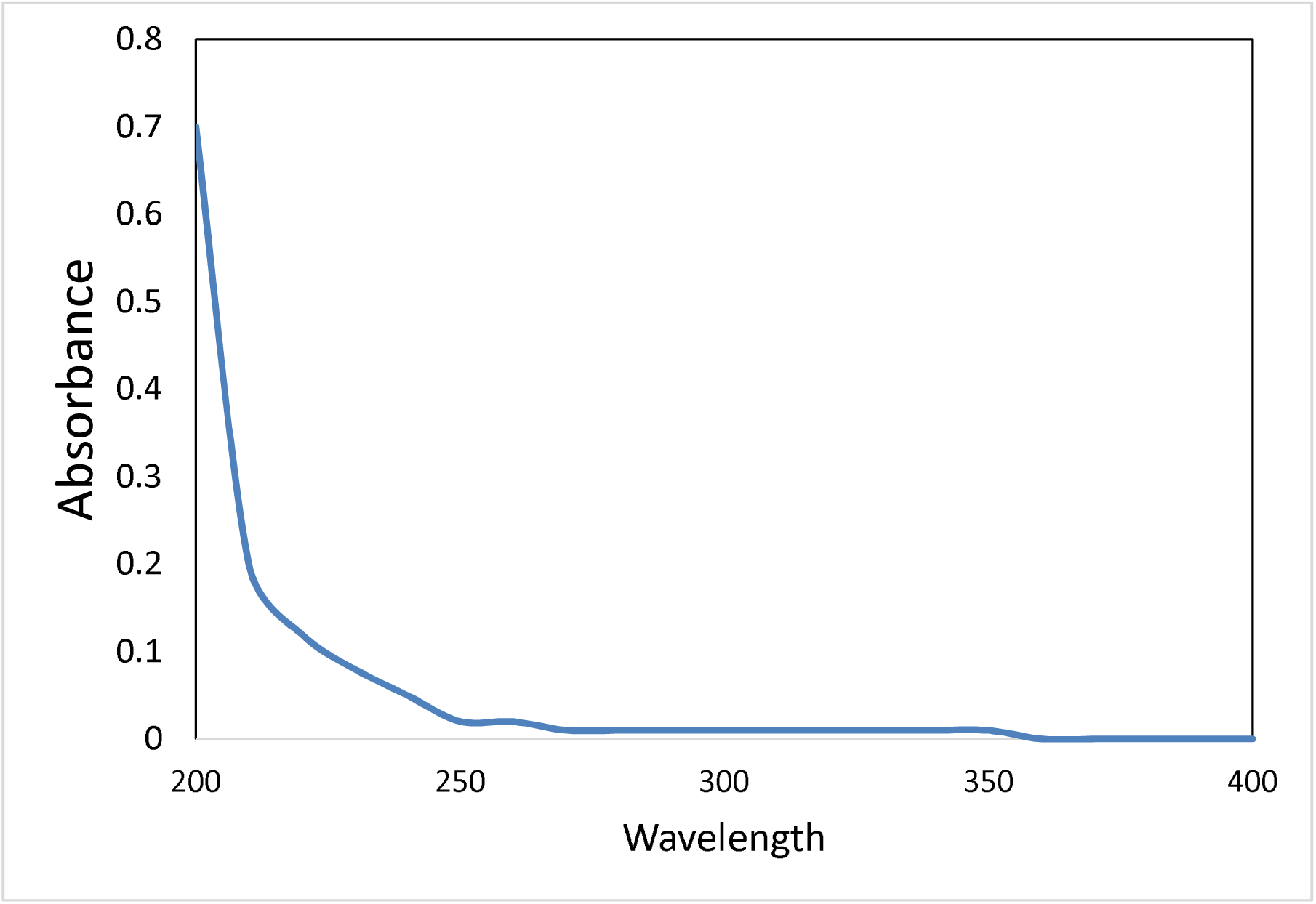
UV-VIS absorbance spectrum of LPS sample shows no absorbance above 215 nm

