## Supplemental Information for "Bacterial lipopolysaccharide induces settlement and metamorphosis in a marine larva"

**This PDF file includes:**

Supplementary text

Figures S1 to S5

Tables S1 to S3

SI References

Supplementary Information Text

Part A: Growth phase analysis of OMVs

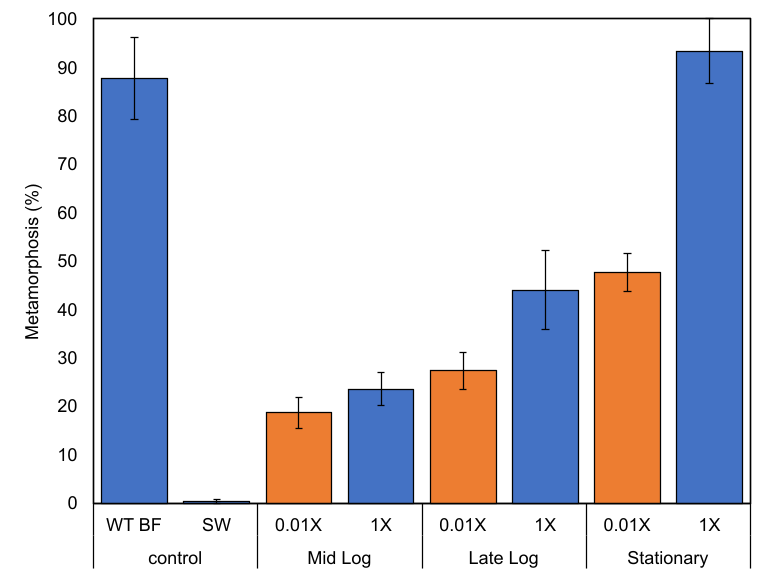

**Figure. S1. Metamorphosis of larvae of *H. elegans* induced by outer membrane vesicles from a culture of *C. lytica* at different growth phases*.*** Metamorphosis of larvae of *H. elegans* when exposed to OMVs from each growth phase (mid log, late log and stationary). 1X is equivalent to the concentration of OMVs found in an inductive 10^8^ cells·ml^-1^ biofilm. 0.01X is a 1/100^th^ dilution. Metamorphosis was counted after 24 h exposure, with filtered seawater serving as a negative control, and a multispecies biofilm serving as a positive control. Letters show significant differences.

Part B. Enzyme treatment of OMVs:

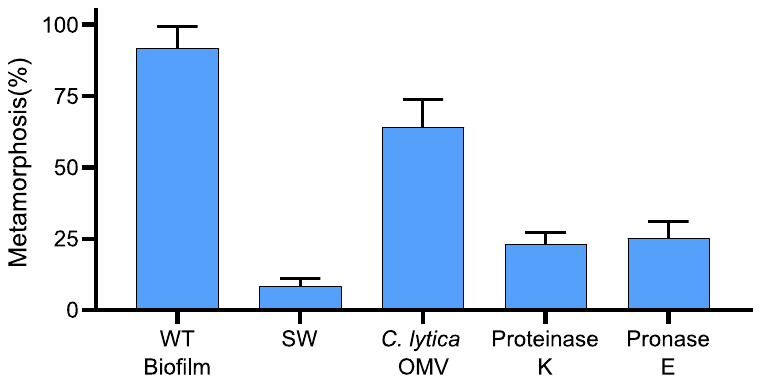

Figure. S2. Proteases induce metamorphosis in larvae of *C. lytica*. Larvae of *H. elegans* were exposed to either Proteinase K (5 U) or Pronase E (5 U). Metamorphosis was counted after 24 h exposure. Filtered seawater (SW) served as a negative control, and a multispecies biofilm and untreated OMVs were used as positive controls.

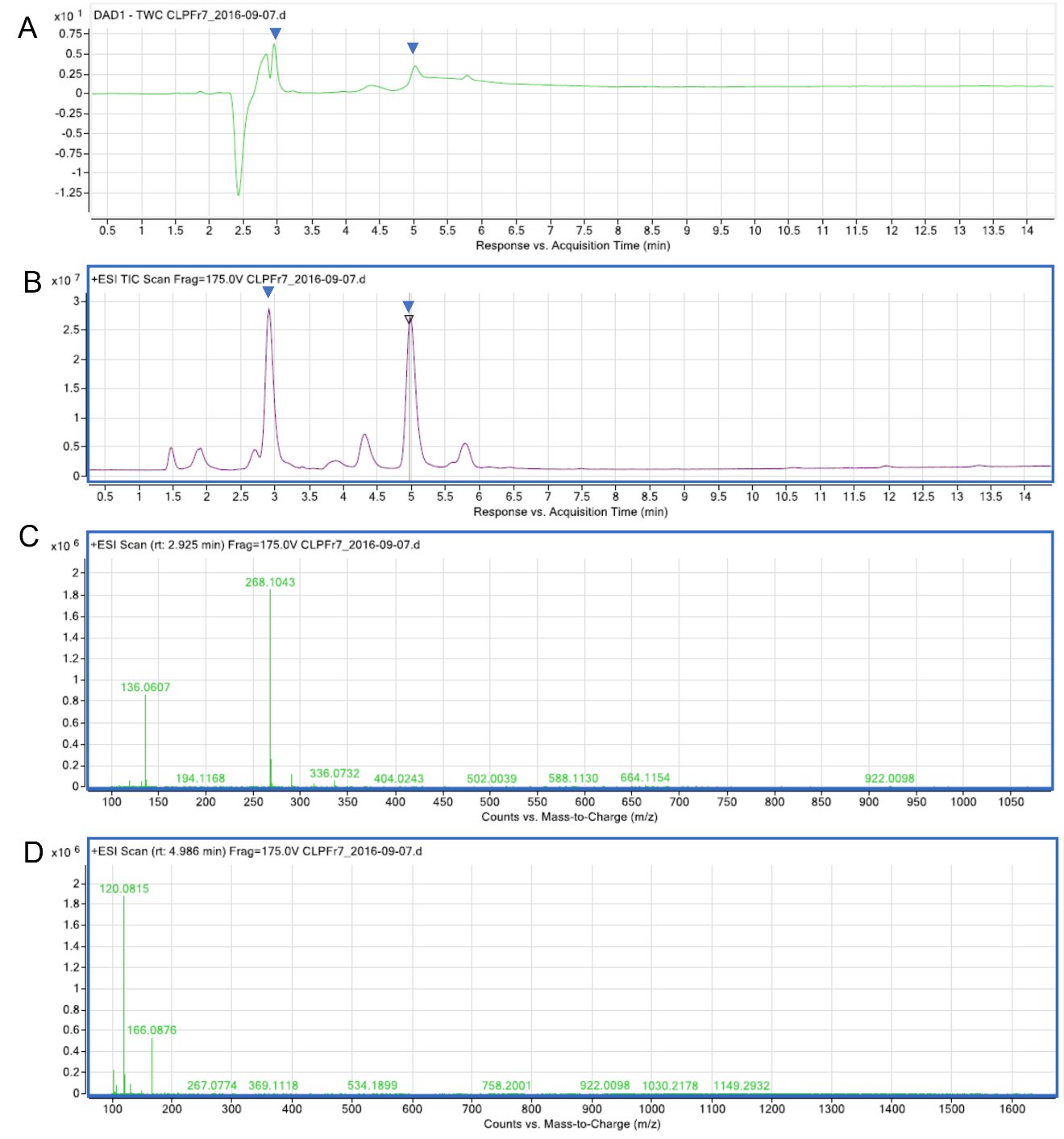
Figure. S3. Spectral confirmation of isolation of adenosine and phenylalanine from most active cell pellet fraction (CLP7). (A) UV-Vis absorbance chromatogram B) Total Ion Chromatogram (TIC) positive mode, Ions were detected within a mass range m/z 100-1100. (C) Extracted ion chromatogram of the active peak at 2.9 min yielded m/z of 268.1043.(D) Extracted ion chromatogram of the active peak at 4.9 min yielded m/z of 123.0459.

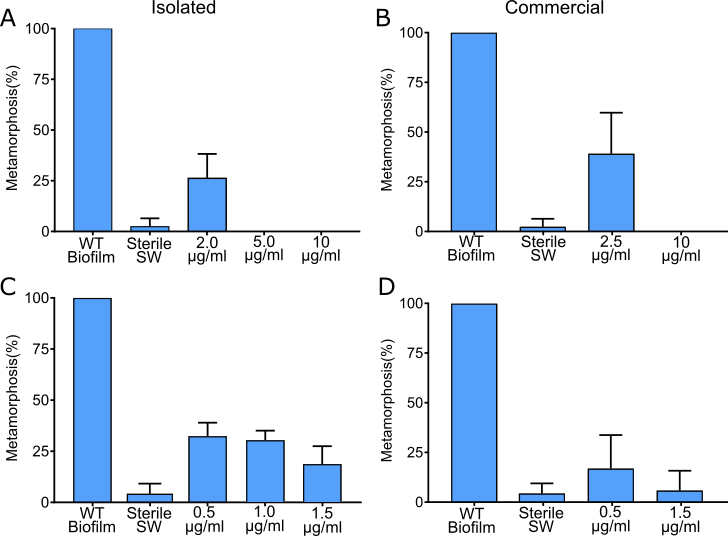

**Figure. S4.** Metamorphosis of larvae of *H. elegans* when exposed to isolated (A) and commercial (C) adenosine and isolated (B) and commercial (D) phenylalanine from Fraction 7 of the cell pellet for 24 hrs. Positive Control: Wild Type (WT) Biofilm. Negative Control: Sterile Seawater (SW).

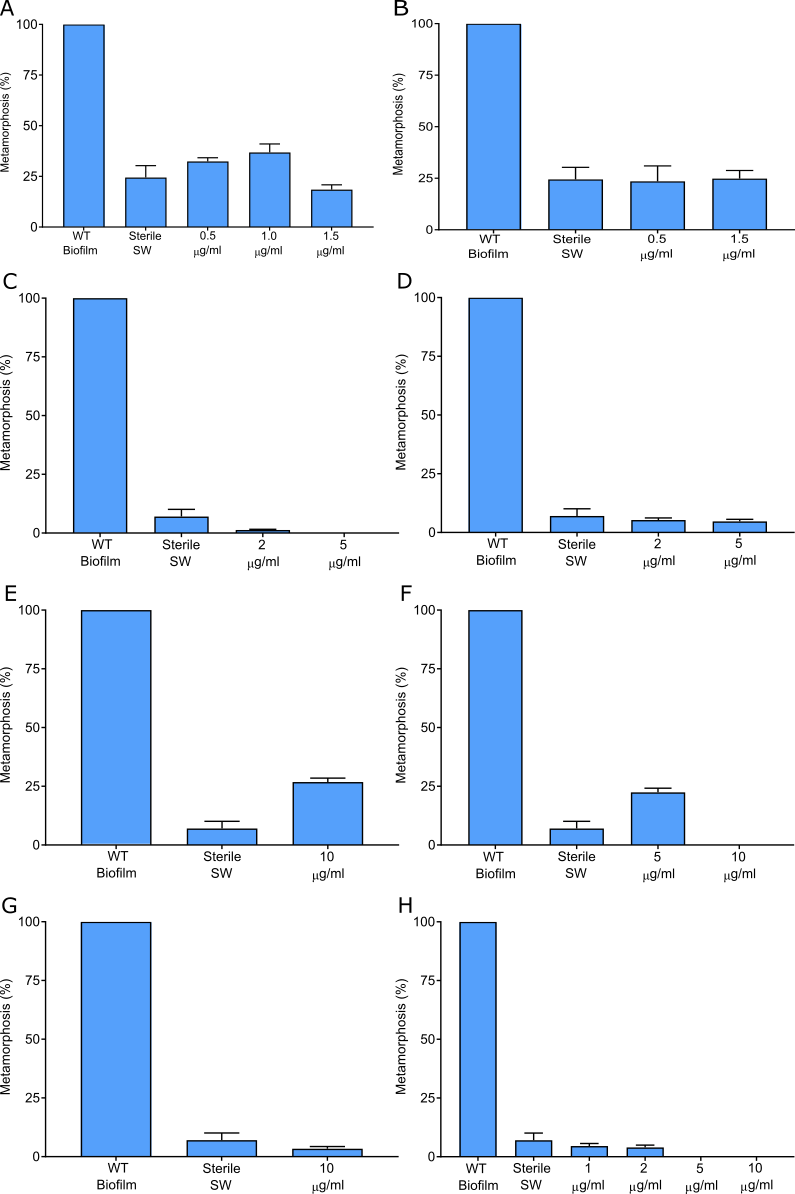

**Figure. S5.** Metamorphosis of larvae of *H. elegans* when exposed to commercial versions of compounds detected in fraction 7 for 24 hrs. A) Adenosine; B) DL-Phenylalanine; C) L-Phenylalanine; D) D-Phenylalanine; E) Niacin; F) Thymidine; G) L=Leucine; H) D-Leucine. Positive Control: Wild Type (WT) Biofilm. Negative Control: Sterile Seawater (SW).

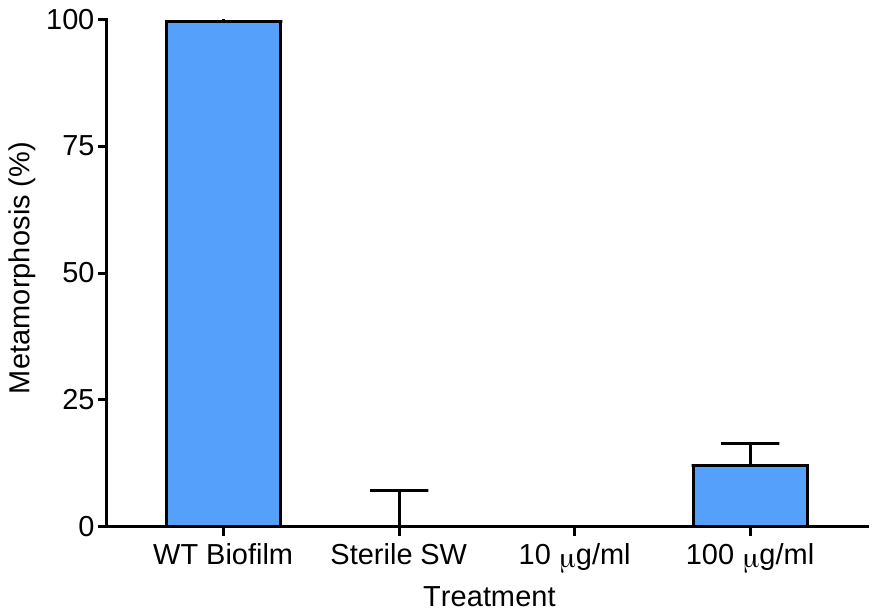

**Figure. S6.** Metamorphosis of larvae of *H. elegans* when exposed to commercial version of the diketopiperizine of phenylalanine and proline (Cyclo(-Phe-Pro) detected in fraction 25 of the supernatant from a *C. lytica* culture for 24 hrs. Positive Control: Wild Type (WT) Biofilm. Negative Control: Sterile Seawater (SW).

Part D. Genome Mining for Secondary Metabolites

Analysis of the genome of *C. lytica* for the presence of secondary metabolite biosynthesis gene clusters revealed a single contig *(ctg1_392)* located between 455,027 - 455,863. A protein blast of this contig, identified it as a phytoene synthase (Table S1). The Minimum Information about a Biosynthetic Gene cluster (MIBiG) failed to match any enzymes with greater than 50% sequence identity (Table S2). Similar low matches were generated using NaPDos (Table S3).

Table S1. NCBI BlastP on ctg1_392 from genome of *C. lytica* generated by AntiSMASH (1)

| Accession | Description | Max Score | Total Score | Query Cover | E value | % Ident |
| --- | --- | --- | --- | --- | --- | --- |
| WP_013620042.1 | phytoene synthase [*Cellulophaga lytica*] | 563 | 563 | 100% | 0.0 | 100.00% |

Table S2. Minimum Information about a Biosynthetic Gene cluster (MIBiG) search results from genome of *C. lytica* generated by AntiSMASH (1)

| MIBiG Protein | Description | MIBiG Cluster | MiBiG Product | % ID | % Coverage | BLAST Score | E-value |
| --- | --- | --- | --- | --- | --- | --- | --- |
| ABB88949.1 | CrtB | BGC0000650 | terpene | 50.0 | 99.6 | 255.0 | 1.2e-67 |
| AAK64298.1 | Phytoene synthase | BGC0000637 | terpene | 37.0 | 100.4 | 168.0 | 1.5e-41 |
| ABD24399.1 | Phytoene synthase | BGC0000644 | terpene | 33.0 | 97.1 | 155.0 | 9.9e-38 |
| AAF65581.1 | Phytoene synthase | BGC0000636 | terpene | 34.0 | 102.5 | 134.0 | 2.4e-31 |
| ABP56869.1 | Phytoene synthase | BGC0001087 | saccharide-terpene | 25.0 | 89.9 | 80.0 | 5.3e-15 |

**Table S3:** Polyketide Synthase (KS) domains within the genome of *C. lytica* generated by NaPDos (2). Search was run with relaxed parameters.

|  | Query ID | Database match ID | % ident. | Align length | e-val | Pathway product |
| --- | --- | --- | --- | --- | --- | --- |
| KS2 | CP009239.1_3_157_15_425 | FabF_Bacillus_FAS | 55 | 409 | 5e^-112^ | fatty acid synthesis |
| KS1 | CP009239.1_4_290_11_391 | FabF_Bacillus_FAS | 26 | 414 | 8e^-24^ | fatty acid synthesis |
| KS3 | CP009239.1_6_112_8_221 | Myca_YP880565_1KSB | 30 | 131 | 2e^-06^ | mycocerosic acid synthase |

Part E: LPS purity checks

Absorbance

| Wavelength | Absorbance |
| --- | --- |
| 200 | 0.7 |
| 210 | 0.2 |
| 220 | 0.12 |
| 230 | 0.08 |
| 240 | 0.05 |
| 250 | 0.02 |
| 260 | 0.02 |
| 270 | 0.01 |
| 280 | 0.01 |
| 290 | 0.01 |
| 300 | 0.01 |
| 310 | 0.01 |
| 320 | 0.01 |
| 330 | 0.01 |
| 340 | 0.01 |
| 350 | 0.01 |
| 360 | 0 |
| 370 | 0 |
| 380 | 0 |
| 390 | 0 |
| 400 | 0 |

**Figure S7.** UV-VIS absorbance spectrum of LPS sample shows no absorbance above 215 nm
